# Highlight Results, Don’t Hide Them: Enhance interpretation, reduce biases and improve reproducibility

**DOI:** 10.1101/2022.10.26.513929

**Authors:** Paul A. Taylor, Richard C. Reynolds, Vince Calhoun, Javier Gonzalez-Castillo, Daniel A. Handwerker, Peter A. Bandettini, Amanda F. Mejia, Gang Chen

## Abstract

Most neuroimaging studies display results that represent only a tiny fraction of the collected data. While it is conventional to present “only the significant results” to the reader, here we suggest that this practice has several negative consequences for both reproducibility and understanding. This practice hides away most of the results of the dataset and leads to problems of selection bias and irreproducibility, both of which have been recognized as major issues in neuroimaging studies recently. Opaque, all-or-nothing thresholding, even if well-intentioned, places undue influence on arbitrary filter values, hinders clear communication of scientific results, wastes data, is antithetical to good scientific practice, and leads to conceptual inconsistencies. It is also inconsistent with the properties of the acquired data and the underlying biology being studied. Instead of presenting only a few statistically significant locations and hiding away the remaining results, we propose that studies should “highlight” the former while also showing as much as possible of the rest. This is distinct from but complementary to utilizing data sharing repositories: the initial presentation of results has an enormous impact on the interpretation of a study. We present practical examples for voxelwise, regionwise and cross-study analyses using publicly available data that was analyzed previously by 70 teams (NARPS; Botvinik-Nezer, et al., 2020), showing that it is possible to balance the goals of displaying a full set of results with providing the reader reasonably concise and “digestible” findings. In particular, the highlighting approach sheds useful light on the kind of variability present among the NARPS teams’ results, which is primarily a varied strength of agreement rather than disagreement. Using a meta-analysis built on the informative “highlighting” approach shows this relative agreement, while one using the standard “hiding” approach does not. We describe how this simple but powerful change in practice---focusing on highlighting results, rather than hiding all but the strongest ones---can help address many large concerns within the field, or at least to provide more complete information about them. We include a list of practical suggestions for results reporting to improve reproducibility, cross-study comparisons and meta-analyses.

**Highlights:**

1. Most studies do not present all results of their analysis, hiding subthreshold ones.
2. Hiding results negatively affects the interpretation and understanding of the study.
3. Neuroimagers should present all results of their study, highlighting key ones.
4. Using the public NARPS data, we show several benefits of the “highlighting” approach.
5. The highlighting approach improves individual studies and meta-analyses.

## Introduction

Reproducibility and replicability have been some of the hottest topics in functional MRI (FMRI), as well as in neuroimaging more generally. Researchers have pointed to various features as underlying problems (or even “crises”) for the field, including: analysis circularity (Kriegeskorte et al., 2009; Vul and Pashler, 2009), publication bias (Jennings and Van Horn, 2012), analysis over-flexibility and the “garden of forking paths” (Carp, 2012; Gelman and Loken, 2014), a lack of sharing analysis pipelines (provenance) and data (Pernet and Poline, 2015; Gorgolewski et al., 2016), *p*-hacking (Wicherts et al., 2016), stringency of cluster threshold levels (Eklund et al., 2016), small sample size and inherently low power in FMRI (Cremers et al., 2017; Turner et al., 2018), having too little data per subject (Nee, 2019), overemphasis on statistics without revealing effect magnitudes (Chen et al., 2017), consideration of stochastic aspects (Tadli and Calhoun, 2022) and artificial dichotomization (Chen et al., 2022b). Some of these issues have prompted widely adopted changes, while others remain points of discussion and controversy.

Here we identify a specific issue that the neuroimaging field should address: the use of all-or-nothing thresholding in published figures and other results reporting, where outputs are only shown for a tiny fraction of the brain, and all other “non-significant” results are entirely hidden from view. Such results are highly sensitive to essentially arbitrary values and often misrepresent the full study data. This form of incomplete results reporting biases interpretations, obscures useful information, and makes meta-analyses---such as properly judging reproducibility---much more difficult. Even though data sharing is increasingly common, such as making group-level datasets publicly available with NeuroVault (Gorgolewski et al., 2015), the present issue is distinct: the initial presentation of figures, tables and other results has an enormous impact on the focus and interpretation of a study, both by the authors and by the readers.

On the face of it, the traditional approach of opaque thresholding has several appealing aspects. FMRI datasets are large (of order 100,000 voxels), so trimming this vast assemblage down to a small set of regions that are related to a task or hypothesis with strong statistical evidence is appealing. By applying thresholds at “standard”, objective levels and following textbook null hypothesis significance testing (NHST), surviving results should all have the same strong statistical evidence. The logic then goes that the output would then contain only those remaining locations/regions worth discussing further, providing useful data reduction: the original input volume is partitioned into a few “active” regions and many “inactive” ones. Furthermore, if the same thresholds are used across studies, then all results should theoretically be on par with each other, allowing for direct comparisons and cross-study analyses. This algorithm has the benefits of being straightforward (apply thresholds, map clusters, and discuss), of being potentially uniform and researcher-independent (just use the same “standard” thresholds, and avoid *p*-hacking) and of leading directly to a simple cross-study reproducibility criterion (do the islands of activity in this study overlap with the islands of that study?).

Unfortunately, this standard approach also has inherent inconsistencies, practical limitations and several undesirable statistical properties, all of which contribute markedly to many of the crises listed above. First, all-or-nothing thresholding enforces the interpretation of brain activation as being ON/OFF like light bulbs. But this picture is fundamentally artificial and inconsistent with basic understanding of neuroimaging and brain behavior: the BOLD response is not a binary process (Gonzalez-Castillo et al., 2012; Orban et al., 2015; Gonzalez-Castillo et al., 2017). Thus, analytic dichotomization inaccurately represents both the data being acquired and the biological processes being studied. Second, there is no universally accepted thresholding approach---the fact that there are several “standard” values and adjustment criteria in use reinforces this point---and the final results can be very sensitive to these criteria (e.g., see Fig. 1 of Cox (2017)). If a cluster ends up having one voxel less than the threshold size, it would be categorized as “below significance” and therefore unreportable. It would be entirely hidden away from the final figures and tables, and never be compared with results of another study for either validation or disagreement. This arbitrary filtering harms the worthy goals of reproducibility and replicability. Third, meta-analyses across studies can produce incorrect interpretations: studies have different power, sample sizes, effect magnitudes and uncertainties, so that even when using the same threshold, what survives will be sensitive to these factors. Due to slightly lower power, one researcher might not see a thresholded cluster that another did and therefore disagree about reproducibility. However, if all effects were reported, they would see the variability is due to differing strength of agreement, rather than to disagreement, which is an important distinction. Finally, all threshold-based outcomes have a fundamentally increased chance of being misleading when statistical power is low (including that some “real” effects remain sub-threshold). FMRI datasets are inherently quite noisy and often of a relatively small size (Cremers et al., 2017; Dumas-Mallet et al., 2017). If increasing the amount of data per individual or increasing the number of individuals in a study reliably increases the number of voxels above a significance threshold (Thyreau et al., 2012; Gonzalez-Castillo et al., 2012; Gonzalez-Castillo et al., 2015), then the statistical threshold does not represent a fundamental property of the data. So, mathematically, opaque thresholding harms reproducibility considerations.

**Figure 1.**
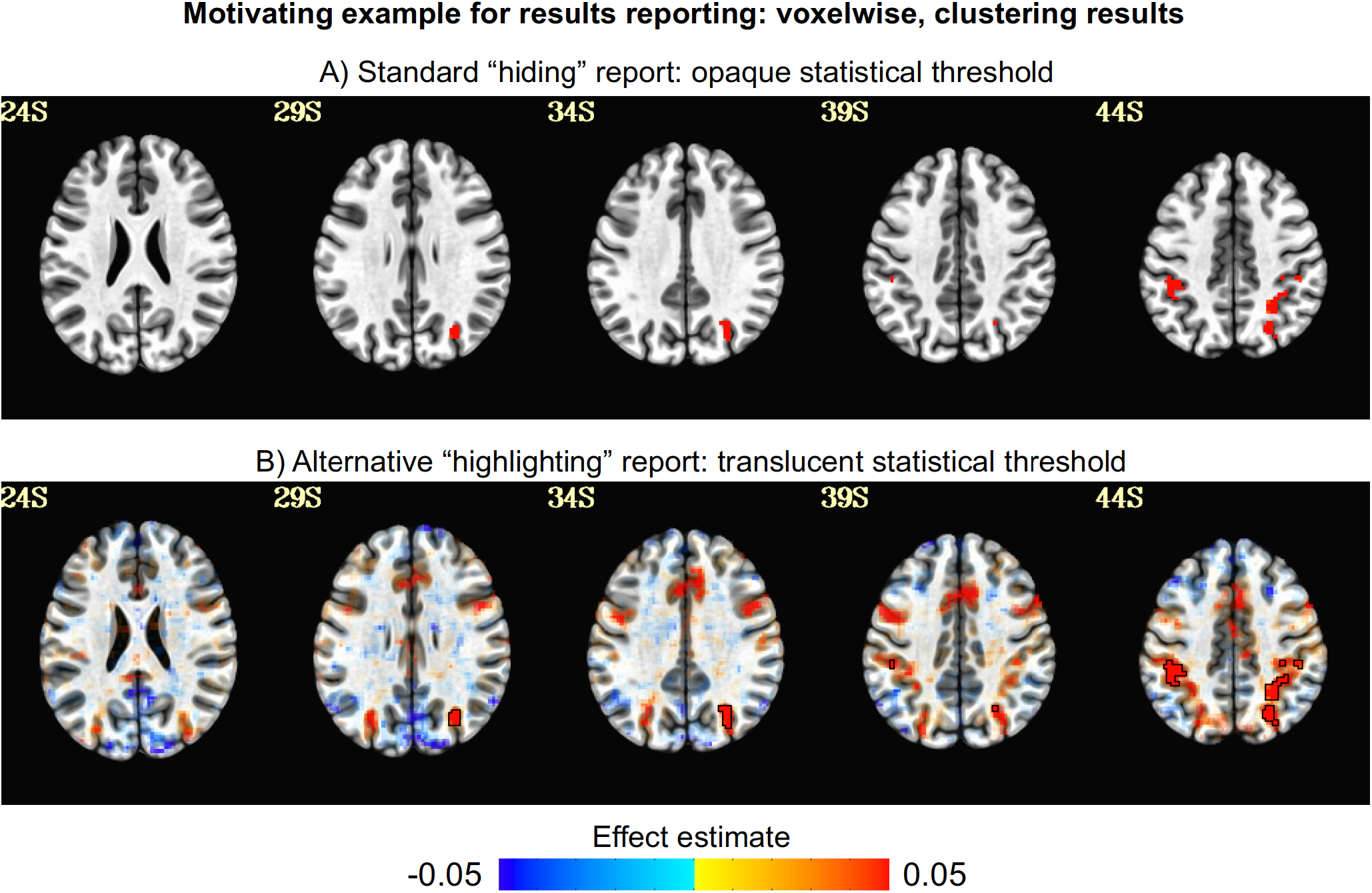
Results reporting matters. Panel A shows five axial slices (left=left) in MNI space from a voxelwise and clustering group analysis for a particular contrast (Flanker task, “incongruent” - “congruent” contrast; see Chen et al., (2022a)) using the current standard approach within the neuroimaging field: only voxels that have passed a threshold of statistical significance are shown, and everything else is hidden away from the reader--even if its value was 99.99% of the selected threshold. Panel B shows the same results with an improved approach, where statistical thresholding is applied with transparency to show more information: suprathreshold results are highlighted by being opaque and outlined with black contours, while sub-threshold locations are visible with decreasing opacity as the voxel’s statistical significance decreases. Which results are a better reflection of the data and results of whole brain modeling? Note that in both panels, the overlay color comes from the effect estimate (BOLD percent signal change), providing useful complementary information rather than just showing the thresholding statistic itself (see Chen et al., 2017). See Appendix A for details on applying translucency in AFNI.

In this paper, we enumerate several issues with the traditional approach of all-or-nothing or “opaque” thresholding.^1^ Some of these have been noted before, as cited below. Briefly, opaque thresholding has the fundamental flaw of hiding scientific outputs: if all voxels in the brain have been modeled, why are most of their results swept under the carpet by arbitrary thresholding and not even shown in images? In most scientific research, it is generally standard practice to report all of the results of fitted models, even those model coefficients that do not rise to statistical significance. It would generally be considered poor scientific practice, or even unethical, to fail to report results of fitted models. In the context of neuroimaging, much useful information may be contained in those excluded results, for the given study as well as for future ones, and should be presented.

We advocate for a methodology in which researchers practice more complete results reporting. This includes using “transparent” thresholding, which both highlights statistically significant regions and also reveals the remaining results. This provides the ability to discuss other scientifically interesting findings. For example, one can leave supra-threshold regions opaque and perhaps even outlined for emphasis, and display sub-threshold regions with decreasing opacity as statistical evidence decreases, rather than zeroing them out (Allen et al., 2012). Thus, all modeling results would be shown, *highlighting* the strongest (so useful data reduction occurs) while still showing all results (and not hiding valuable information).

Here, we illustrate our proposed approach of transparent thresholding using real data and existing software implementations (e.g., in AFNI, Cox (1996)), for voxelwise, region-wise and cross-study analyses. We utilize the publicly available FMRI data from the NARPS Project (Botvinick-Nezer, et al., 2020), in which 70 teams independently processed the same data and the results were studied for variability. We also point to other useful techniques for data presentation, such as displaying effect estimates and model evaluations, which are also in line with more complete results reporting.^2^ Finally, we discuss the numerous mathematical, biological and practical issues with traditional thresholding, as well as how the proposed “highlighting” approach resolves many of those difficulties. This small and essentially cost-free adjustment for the field---moving from a “hiding” mindset of results reporting to a “highlighting” one---would have many benefits for reproducibility, verification, quality control, meta-analyses and more issues that have been raised in the field.

## Background: A motivating example

Traditional results reporting in neuroimaging uses opaque, all-or-nothing thresholds, and only elements that pass one or more thresholds are reported and displayed. This means that the statistical outcomes throughout the vast majority of the dataset are entirely ignored in the presentation and immediate interpretation of the study. Figure 1A shows an example of presenting group-level thresholded results in the standard all-or-nothing thresholding way (e.g., voxelwise thresholding at *p*=0.001 and clusterwise thresholding based on a familywise error rate FWE=5%). Supra-threshold results are labeled “active”, and sub-threshold results are hidden away as if absolutely nothing happened there. Is that really an accurate reflection of the brain behavior, acquired data and statistical modeling?

Now consider Figure 1B, which shows the exact same dataset but allows subthreshold results to be observed alongside the supra-threshold ones using a visualization method first suggested^3^ for voxelwise reporting by Allen et al. (2012), but not widely adopted. Here, voxels whose values are above threshold are opaque and outlined, and the remaining ones are shown with decreasing opacity as statistical significance decreases. One can observe how much valuable information has been lost in Panel A—it is certainly not true that nothing of interest occurred in the brain outside of the few supra-threshold clusters, even if the statistical evidence is weaker. Consider the presence of bilateral symmetry in the patterns of Panel B that were essentially hidden in Panel A, as well as the recognition of spatial patterns of several known functional networks. Panel B presents more informative results, using opacity and outlines to highlight a specific subset, while Panel A only presents a tiny proportion of results and hides the vast majority of the analytic information.

The “highlighting, not hiding” (or simply “highlighting”) approach can be applied much more generally to other scenarios of results reporting and model validation. We describe several more cases below.

## Methods: Practical examples for improving results reporting

We provide three practical demonstrations of the proposed “highlighting” methodology vs the standard “hiding” approach, using real FMRI data. These include: 1) voxelwise group analysis; 2) regionwise group analysis; and 3) a cross-study (or meta-) analysis. Details on how translucency is calculated and applied within the AFNI visualization are provided in Appendix A.

### Background: public dataset used

We utilize FMRI data from the recent NARPS project by Botvinik-Nezer, et al. (2020), which was aimed at studying the variability in findings when a single data collection was processed, analyzed, and reported by 70 teams. Participating teams were given data from two groups (“equal indifference” and “equal range”) of 54 subjects each; each subject had one T1w structural and one task-based FMRI dataset acquired at 3T (for further details, see Botvinik-Nezer, et al. (2020) and https://www.narps.info/analysis.html#protocol). The task was a mixed gambling paradigm, with each stimulus event comprising a pair of potential gain and loss values, to which the subject was supposed to respond quickly in one of four levels (strongly/weakly accept/reject).

In the main NARPS project, the organizers posed nine hypotheses: each referring to a specific group, to one specific gain/loss response, and to one of three anatomical brain regions.^4^ Participating teams were asked to “report a binary decision (yes/no) based on a whole-brain correction analysis” for each. Teams were allowed to use any software, processing stream, regression/modeling options (e.g., whether to use or ignore response levels, as well as cases where no response was given; how to view the duration of an event; and whether to orthogonalize or modulate events), whole brain statistical correction approach and even region of interest (ROI) definitions (no atlas was provided).

Contributions from 70 teams were submitted, of which 65 were used in the final pooling of thresholded results. Of the nine hypotheses, 94% of teams reported “no” for hypotheses #7, 8 and 9; 84% reported “yes” for #5; approx. 78% reported “no” for #2-3; 67% reported “no” for #4 and 6; and the lowest agreement was 63% reporting “no” in #1.^5^ Thus, 6 out of 9 hypotheses had approx. 80% or higher agreement in thresholded/binarized outcomes across teams, from (pre-quality control) group sizes of just 54 subjects each.

### Processing for voxelwise and regionwise analyses

Processing of the NARPS data was performed with AFNI (Cox, 1996) and FreeSurfer (Fischl and Dale, 2000), for both voxelwise and region-based analyses. Both subject- and group-level processing and modeling are described below briefly, and full code is provided https://github.com/afni/apaper_highlight_narps. The voxelwise group results used in Figs. 2-3 and the datatable group results used in Figs. 4-5 are in NeuroVault Collection 13104. See the original NARPS paper for links to the individual team results used in Figs. 7-9 and Supplementary Info.

**Figure 2.**
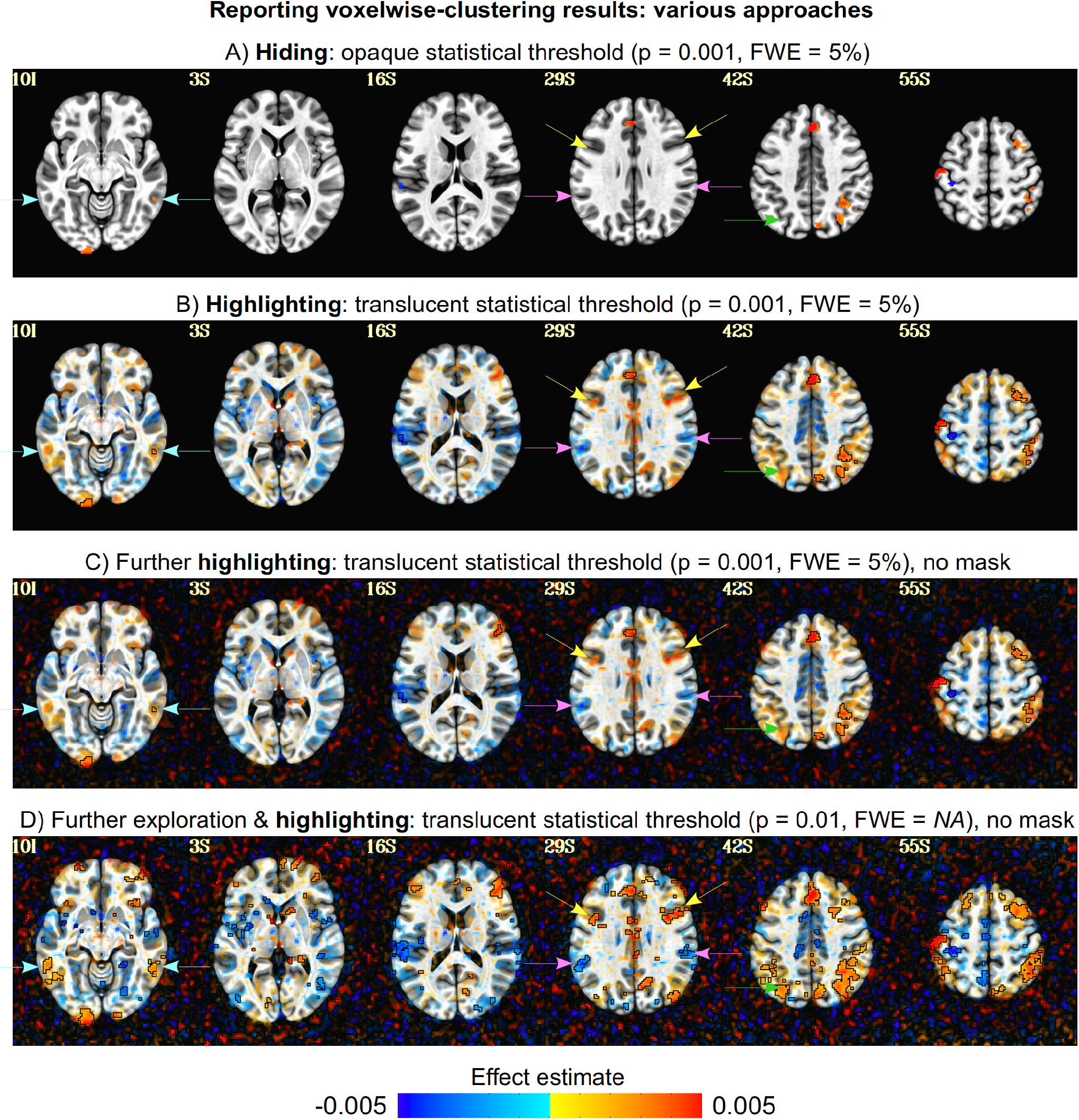
A comparison of group-level effect estimation for voxelwise results (parametric gain condition, equal range group, used in Hyp #2 and 4) using the “hiding” and “highlighting” styles (axial slices: left=left), with effect units of BOLD % signal change per dollar. Panel A shows the standard opaque thresholding, where most of the brain contains no information for helping to interpret supra-threshold regions or to understand modeling results there. Panel B shows the more informative approach, of highlighting supra-threshold regions and also showing the surrounding context translucently. Panel C uses the same thresholding as Panel B but without brainmasking applied, which allows for potential artifacts or alignment issues to be observed. Panel D shows another example using the “highlighting” style, at a lower threshold and minimal clustering (a nominal threshold of 5 voxels), for further exploration and comparison. The arrows show several cases where opaque thresholding might leave questions about how to interpret a cluster (e.g., a small one without bilateral symmetry), but with transparent thresholding more context is provided (such as bilateral partners, even if with statistically weaker evidence) for a richer picture.

**Figure 3.**
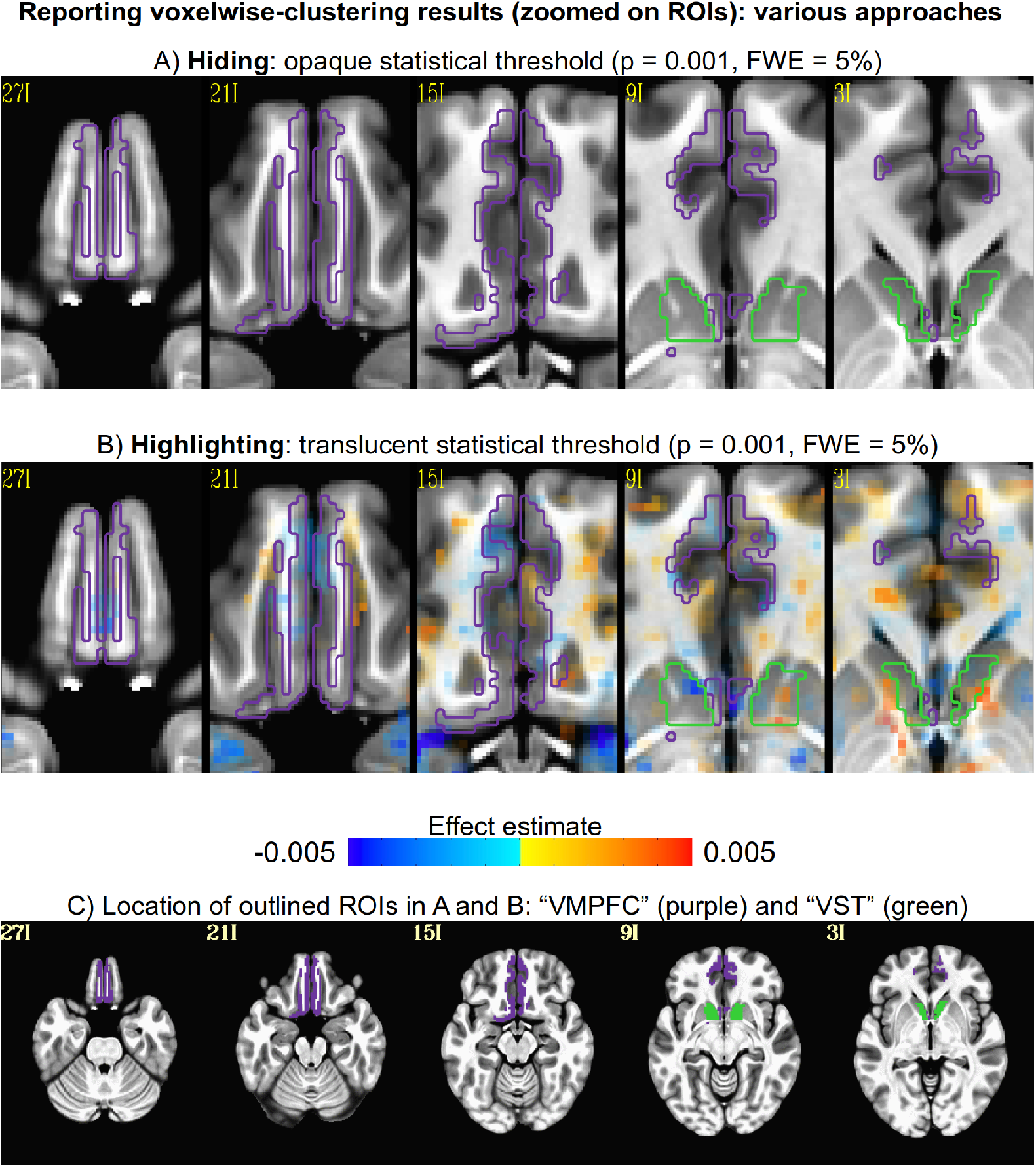
A comparison of voxelwise group-level results, corresponding to those in Fig. 2A-B but zoomed in on the specific regions of the NARPS Hyp #2 (“VMPFC”, purple outline) and 4 (“VST”, green outline) (axial slices: left=left). As in Fig. 2A, opaque thresholding provides no information about results in these regions, if no voxels survived the dual-thresholding. In contrast, and mirroring Fig. 2B, the transparent thresholding provides information in all cases; even if no voxels were colored here, that would still provide the information that those locations have extremely weak statistical evidence, rather than the less informative fact of just being somewhere below threshold. Panel C shows the locations of the VMPFC (purple) and VST (green) on the MNI template.

**Figure 4.**
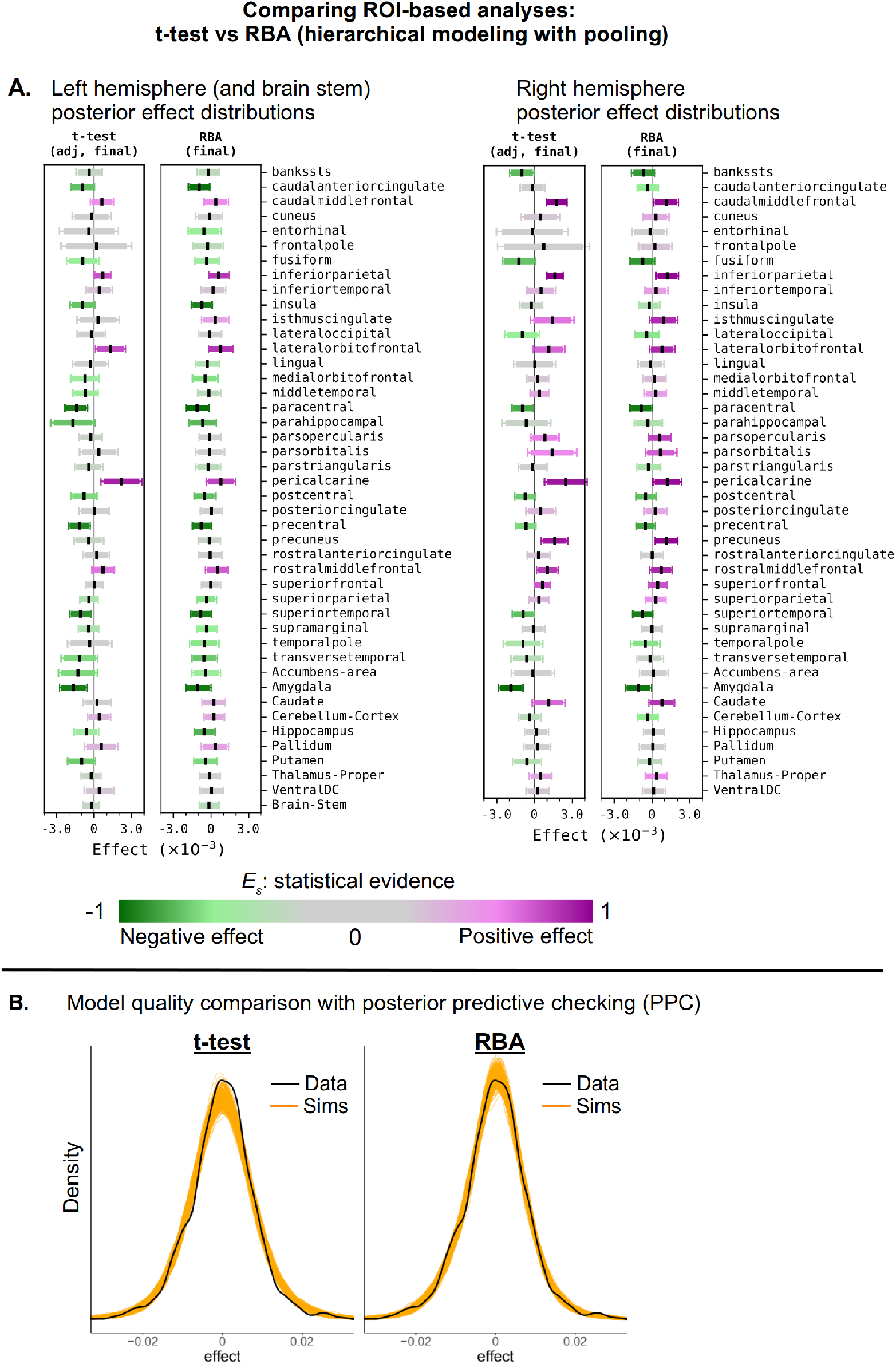
A comparison of group-level effect estimation for ROIs with the 87-region Desikan-Killiany atlas using two modeling approaches. Panel A: For each ROI, boxplots show the point estimates and the uncertainty ranges: the t-test result (left column) and the hierarchical modeling result (right column) with RBA. For each boxplot, the black dot shows the median, the box contains the 90% uncertainty interval, and the whiskers show the 95% interval. Here, the color shows the extent of statistical evidence E_s_ for the effect (not the value of the effect itself, as in the brain map images): a value close to 1 (or -1) indicates strong evidence for a positive (or negative) effect. The E_s_ coloration for the t-tests is based on the FDR adjusted q-value, while the boxplot interval reflects the original statistic. In most cases, the impact of partial pooling can be seen with the median of hierarchical modeling closer to center than its counterpart from the t-test, as well as the uncertainty interval shrinking. Interestingly, the strength of statistical evidence is quite similar between the adjusted t-test and RBA values. Panel B: Posterior predictive checking (PPC) provides a method for comparing the quality of the model results to the original data via Monte Carlo simulations, using the effect estimate information. Here, the hierarchical RBA method provides a much better quality than the t-test results, reflecting the relative benefit of the former’s partial pooling.

**Figure 5.**
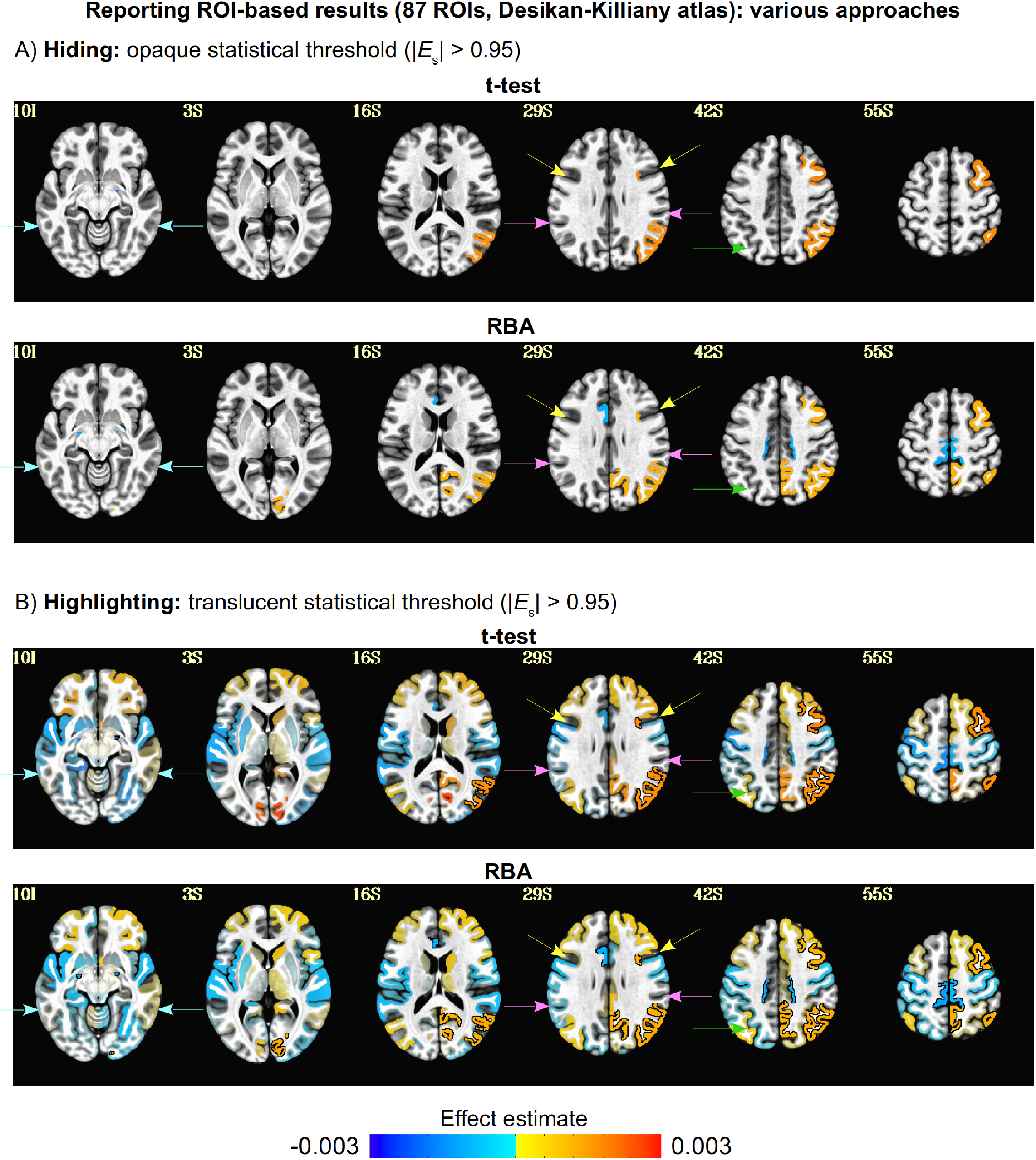
A comparison of group-level effect estimation for ROI-based results with the 87-region Desikan-Killiany atlas using the traditional style of “hiding” results and the newer “highlighting” style (axial slices: left=left). Results of two different statistical methods are shown (see corresponding boxplots in Fig. 4A): t-test results after FDR adjustment, and the hierarchical modeling output from RBA. Here, panel A displays each of the model results using standard opaque thresholding, while panel B displays the same results using transparent thresholding. For reference, the arrows show the same locations as in Fig. 2. Here, as well as in voxelwise plots, the transparent thresholding shows a more informative picture of results. We note that while these results are generally consistent with voxelwise results for locations of high significance and sign of effect, the range for the magnitude of effects in the ROIs is typically narrower, due to averaging over a larger and more heterogeneous volume.

First, the anatomical volume was processed as follows. FreeSurfer’s recon-all was run on each subject’s anatomical volume to generate whole brain parcellations. In particular, we utilized the Desikan-Killiany atlas (Desikan et al., 2006) in different ways for the voxelwise and regionwise analyses (described in respective sections, below). AFNI’s @SSwarper was used to perform skullstripping (SS) on the anatomical scan, as well as to estimate nonlinear transformations (warps) to standard MNI-2009c space.

AFNI’s timing_tool.py was used to create timing files from each subject’s events.tsv file. We chose to use gain and loss values as continuous amplitude modulators, rather than to use discrete response classes (weak/strong by accept/reject) as separate conditions. Therefore, all events with responses were collapsed into a single “response” (Resp) condition with 2 modulators (gain and loss), where the event duration was given by the response time. Events without any response were coded as a separate “no response” (NoResp) condition, with events lasting the full 4 s duration.

AFNI’s afni_proc.py was used to create the full FMRI processing pipeline through regression modeling for each of the voxelwise and regionwise analyses. For the former a blur of FWHM=4 mm (twice the input EPI voxel dimensions) was included, and for the latter no blurring was performed, since the time series would be averaged within each ROI and the signals should not be spread across region boundaries. All other processing options were the same.

Results from @SSwarper (the skullstripped anatomical and nonlinear warps), timing_tool.py (stimulus timing files) and FreeSurfer’s recon-all (anatomical parcellations) were all included within the afni_proc.py-generated script. EPI-anatomical alignment was calculated using the lpc+ZZ cost function (Saad et al., 2009), also checking for possible left-right flips (Glen et al., 2020). For ROI-based analyses, spatial blurring was not applied. Fast ANATICOR (Jo, et al., 2010) was included as part of motion reduction. Motion censoring was included for time points with Enorm (Euclidean norm) > 0.3 mm or an outlier fraction >5% within a whole brain mask. Resp events used “gain” and “loss” as amplitude modulators (demeaned, to keep the modulation regressors orthogonal to the main response regressor), while using the response time for the duration. During processing each time series was scaled to local BOLD percent signal change, so that the main Resp coefficients had these units, and the Resp modulation coefficients for this mixed gambling task had units of BOLD % signal change per dollar. NoResp events were encoded without any modulators, while using the full 4 s duration. Voxel-level temporal structure of ARMA(1,1) in the residuals of the subject-level time series regression model was estimated and accounted for using 3dREMLfit.

afni_proc.py’s QC HTML was used for detailed quality control evaluation of each subject’s processing. In conjunction with the focus of this study’s topic, we note that the QC report’s volumetric images of statistics and correlation patterns are purposefully presented with transparent thresholds for full quality evaluation. Finally, gen_ss_review_table.py was used to exclude subjects with: censor fraction >10% of initial volumes; large average censored motion, mean(Enorm) > 0.1; or large maximum censored displacement, max(Enorm) > 8.

### Voxelwise analysis

Following the original NARPS formulation, we performed whole brain voxelwise analysis but also focused on three specific regions of interest. We utilized the Desikan-Killiany atlas to provide the locations of the three ROIs from the stated NARPS hypotheses: the ventral striatum (VST), ventromedial prefrontal cortex (VMPFC) and amygdala, each without separate left/right consideration. Only the latter of those regions is labeled directly in the Desikan-Killiany atlas from FreeSurfer. Therefore, here we have used the following parcellation ROIs to closely match those regions: {Left,Right}-Accumbens-area for VST; ctx-{l,r}h-medialorbitofrontal for VMPFC; and {Left,Right}-Amygdala for amygdala.

The nine hypotheses of interest in the NARPS project were explicitly 1-sided. However, for the whole brain analysis here we used basic 2-sided t-tests, because that is how most FMRI hypotheses are or should generally be formulated (Chen et al., 2019a). We also note that NARPS Hyps #5-6 essentially form a 2-sided pair with Hyp #7-8 at the whole brain level. This may lead to a final outcome difference in statistical level, compared to teams that used uncorrected 1-sided testing.

For this voxelwise example (and for the regionwise one, below), we analyze and discuss the data that apply to Hyp #2 and 4: the “parametric effect of gain” in the “equal range” group. The cluster threshold for the voxelwise analysis was calculated nonparametrically for the blurred data using AFNI’s 3dttest++ with the “-Clustsim” option (Cox et al. 2017a; Cox et al. 2017b), within a group-intersection mask. We first present whole brain results (as the present work is a general discussion of results presenting), and then zoom in on the specific regions associated with the hypotheses: the VMPFC (Hyp #2) and ventral striatum (Hyp #4).

### Regionwise (ROI-based) analysis

We investigated ROI-based analyses using two separate parcellations that of differing coarseness. First, we used all 87 ROI-like GM regions from FreeSurfer’s Desikan-Killiany atlas, which were generated for each subject as noted above. These were input as “follower” datasets within the initial afni_proc.py, so that they were transformed to the final EPI grids in MNI space for each subject. Second, we used the 360 ROI Glasser atlas (Glasser et al., 2016), which is already defined in MNI space.

We also used this example to illustrate how the highlighting approach can be applied with both a conventional t-test analysis and in a Bayesian analysis. Furthermore, we show how the latter has several advantages for more informative results reporting. Firstly, the conventional t-test is performed as a massively univariate analysis, and therefore requires some kind of statistical adjustment for the multiple comparisons performed (FDR, Bonferroni, etc.). The Bayesian modeling implemented here removes this level of fairly arbitrary statistical adjustment (“arbitrary” since the different post hoc approaches provide very different results) through having a single hierarchical model structure.

Secondly, Bayesian modeling facilitates model validation, an important part of results reporting, with posterior predictive checking. As a last point for investigating parallel analysis approaches, in viewing the two sets of results themselves, we see the benefits of the highlighting approach, since it provides a more informative comparison and one that is less sensitive to small differences.

For each subject, the unblurred preprocessed results were averaged within each of the atlas ROIs. For each atlas, the two statistical approaches were applied in parallel, for comparison. First, standard 2-sided t-tests were run for each region in a “massively univariate” fashion. From the many available approaches to then account for multiple comparisons, standard false discovery rate (FDR) adjustment was applied to the statistical values, since Bonferroni correction would be overly strict (and such corrections can be damaging for reproducibility and meta-analysis accuracy, on their own).

Second, Bayesian multilevel modeling with AFNI’s RBA program for “region-based analysis” (Chen et al., 2019b) was used for a “region-based analysis” that adopts hierarchical modeling and partial pooling under the normal distribution assumption of cross-region effects, so that one single model can be used to efficiently calibrate information across space without further need for multiple comparisons adjustment (Gelman et al., 2012; Chen et al., 2019b). We demonstrate using model quality evaluation to compare the t-test and RBA analysis approaches, and also implement a leave-one-out cross-validation (LOO-CV) technique for further comparison (Vehtari et al., 2017).

We note that the t-test and Bayesian multilevel modeling methods provide similar but slightly different “standardized” statistical quantities for comparison. The former’s t-statistic can be converted to the common *p*-value, while the latter’s statistic can be reduced to a quantity *P*^*+*^, which is defined as the posterior probability of the effect being positive (calculable as the area under the posterior predictive curve for effect>0). While the interval of *P*^*+*^ is [0, 1], unlike the *p*-value its value carries an inherent directionality of effect: values close to 1 signify strong evidence for a positive effect, and those close to 0 signify strong evidence for a negative effect. Therefore, to be able to compare the statistical outcomes on a similar scale, the *p* and *P*^*+*^ variables were mapped to a common “statistical evidence” quantity *E*_s_ with a range of [-1, 1] by construction, as follows:

- *E*_s_ = (1 - *p*) for a positive effect, and *E*_s_ = -(1 - *p*) for a negative effect;
- *E*_s_ = 2*(*P*^*+*^ - 0.5) for all *P*^*+*^.

The interpretation of *E*_s_ is then the same across the tests: values close to -1 indicate strong statistical evidence for a *negative* effect; values close to +1 indicate strong statistical evidence for a *positive* effect; and values close to 0 indicate a *lack* of statistical evidence for any effect.

### Cross-study (“cross-team”) analysis

As a final means of assessing the benefits of transparent thresholding, we downloaded the group-level statistics maps submitted by the teams who participated in the NARPS project. These were available from NeuroVault (Gorgolewski et al., 2015). We excluded the same 6 teams’ results that the original NARPS project excluded from their unthresholded analysis. Additionally, the Neurovault uploads from three other teams (IZ20, P5F3, Q6O0) contained files that were ambiguously named and which could not easily be connected to particular hypotheses. Therefore, these were excluded, and results from 61 teams were used here.

We note that the uploaded statistical maps could be t- or z-values, so there is a mix across all participating teams. For simplicity, we treat these as approximately equivalent, under the assumption that most teams’ final group sizes were around 50 subjects.

We investigated the similarity and variability across these teams’ results. Here, we present an example exploring the “parametric effect of gain” in the “equal indifference” group, which was used for NARPS Hyp #1 and 3. First, each of the “hiding” and “highlighting” methodologies were applied for presenting the cross-team results at the whole brain scale, and we compared the relative interpretations of these representations. These included both visualization of the data and the calculation of cross-team similarity matrices for quantitative characterization. Then, since the NARPS hypotheses focused on specific regions (VMPFC and VST for #1 and #3, respectively), we explored a comparison of highlighting and hiding approaches in the subregions encompassing those ROIs. The Supplementary Information contains the same analyses for the remaining NARPS hypotheses.

## Results

In this section, we present voxelwise, regionwise and cross-study results. In each case, we compare the presentation of the exact same modeling results in two separate ways: with standard all-or-nothing thresholding, and with the proposed transparent thresholding. We highlight the benefits of the latter’s “highlighting” style, demonstrating how it generally contains more useful information to guide interpretation and less sensitivity to threshold values.

Furthermore, we point out additional ways to apply the “highlighting” approach to present more full results. For the voxelwise analysis, we show how summarizing cluster results with “peak coordinate” tables hides too much information, while relative overlap tables provide a more complete “highlighting” approach. For regionwise analysis, we show how hierarchical Bayesian methods have similarities to standard massively univariate approaches, but also important advantages: they reduce threshold sensitivity by removing the need for arbitrary post hoc adjustments, and they provide more complete modeling information with built-in posterior validation methods. Finally, in the cross-study analysis we show how the highlighting approach allows for a more complete interpretation of variability, differentiating between conflicting results or simply agreement with different magnitudes.

### Voxelwise results

After preprocessing and QC, there were 47 subjects included in the “equal range” group and 52 in “equal indifference” for group-level analyses. Here, we present the results applicable to Hyp #2 and 4: the “parametric effect of gain” in the “equal range” group. A standard voxelwise analysis with clustering was performed, using a voxelwise threshold of *p*=0.001 and cluster-level threshold of FWE=5% (which was 28 voxels here). There were 15 regions above the dual threshold, and the standard style of all-or-nothing reporting for these results is shown in Fig. 2A. Note that the presented results leave many interpretational questions: for small ROIs, should we be dubious if they are not bilateral (see cyan arrows)? Even if some clusters are bilateral, what if they are borderline-small and/or oddly shaped (cyan arrows)? What should be made of the sizable asymmetry between some large and small regions (green arrows)?

These questions are quickly resolved by simply presenting the same results with transparent thresholding, as shown in Fig. 2B. Several locations now show bilateral symmetry, if at slightly lower significance (see cyan and pink arrows). While this is not necessarily “proof” of underlying activity (many tasks would be expected to be lateralized or asymmetric), there are many cases where such spatial patterns are of great physiological interest. The presence of odd shapes in the thresholded results may also be seen to be an artifact of the thresholding---the clusters are just the peak of a more smoothly-valued region (pink arrows). A small cluster in slice Z=42S (see green arrow) appears to be part of a larger region of lower activation that more closely parallels symmetric activation in the other hemisphere. Additionally, new locations of possible interest appear, which might be observed in a study with larger numbers of trials or subjects, with an improved model, or with a slightly different paradigm. Consider, for example, the bilateral positive-effect regions in the inferior frontal gyri in slice Z=29S (yellow arrows).

Note that it is neither a necessary nor a universal property that isolated regions uniformly gain slightly sub-threshold symmetric partners with this new approach. What is important is that the researchers and readers are aware of this “extra” information immediately, whether it is the presence of symmetric near-activation, or symmetric activation of the opposite sign, or a lack of strong evidence for activation. At the same time, the highest-significance regions are still highlighted, so there is no loss of information compared with opaque thresholding.

Fig. 2C uses the same thresholding as Panel B, but without brainmasking applied. This allows for the visualization and interpretation of artifacts that might affect within-brain results, such as ghosting or coil issues, as well as of potential influences of bordering structures (such as dura or skull). Additionally, one can verify the quality of alignment. The default FMRI processing within afni_proc.py calculates masks for the data, but does not apply them, so that the full field of view can be observed for these reasons (e.g., in the HTML report images; see the Discussion). The benefits of this quality control (QC) aspect can be further appreciated when observing several of the NARPS teams results in Fig. 8.

Fig. 2D shows that this highlighting visualization is flexible: the same results can be viewed at various thresholds, with minimal or no clustering applied (a nominal cluster threshold of 5 voxels is applied in Panel D, for noise reduction). Our proposed transparent results reporting approach is less sensitive to thresholding, so the results reporting is more stable and the change in visualization and interpretation is relatively small, even when adjusting the voxelwise *p* threshold by an order of magnitude. This flexibility may be useful for data exploration or visual comparison of datasets at a lower level of thresholding. The items that are highlighted might differ slightly, but the results are still reported with more stability and completeness.

Since the original NARPS paper looked at specific ROIs for each hypothesis, in Fig. 3 we “zoom in” on the results shown in Fig. 2 for the regions specifically investigated in Hyp. #2 and 4. Fig. 3A provides no information at all about the results that overlapped these regions, other than no evidence appears to have supra-threshold statistical significance there. We are unable to assess whether there were any patterns in the modeled results, and we are not even certain that the final brain mask included this region (e.g., it might have been excluded due to signal dropout and low SNR). However, the “highlighting” methodology in Panel B shows much more information about the modeling. In this particular example no clear pattern emerges, but at least the reader has valuable information about the modeling outcomes there.

One additional aspect is worth noting. A common practice in voxelwise results reporting is to summarize each cluster’s location using its peak statistical voxel (or center of mass), providing a single coordinate and/or the name of an atlas region there. In theory this is a useful way to list out important locations, but in practice this does not accurately summarize clusters: they generally cover multiple sulci and gyri and encompass multiple ROI boundaries. The location of peak statistical values are unstable---nor are they necessarily located at the location of peak effect magnitude---and no single coordinate accurately summarizes complicated cluster shapes. Table 1 displays representative tables of this standard approach for the suprathreshold clusters shown in Fig. 2A-B, and it shows the arbitrariness inherent in this approach: atlas association can vary depending on the summary location chosen (4 out of 15 clusters). Reducing the entire cluster extent to a single “summary” voxel becomes a further *hiding* approach in results reporting, losing important information about the determined extent of the cluster (even though the cluster methodology *specifically* leverages spatial extent for statistical reliability).

**Table 1.**
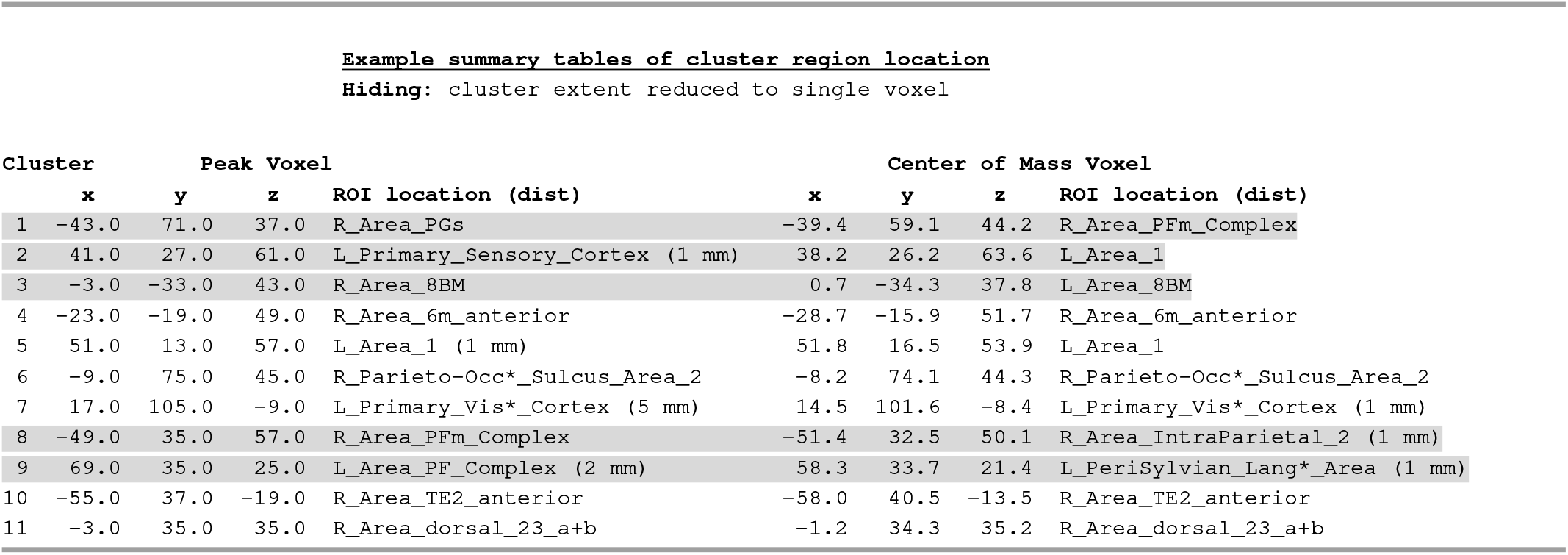
Examples of standard tabular reporting and summarizing of voxelwise group results after clustering (with the Glasser atlas for reference location; coordinates are in RAI Dicom notation). Clusters are typically summarized by just the location of their peak voxel (left side), or occasionally by their center of mass (right side). In either case, the full extent of the ROI is shrunk to a single location and region, even though it often overlaps with multiple regions. The parenthetical “dist” refers to the distance to the nearest atlas ROI when the reference voxel is not contained in one. Cases are highlighted where the choice of peak or central voxel would lead to different ROI associations (5 out of 11). Reducing cluster extent to a single voxel is a “hiding” approach that loses much useful information and is sensitive to an arbitrary choice of meaningful voxel. This can be contrasted with the more full results reporting of Table 2, which highlights the relative overlap of each cluster.

Alternatively, one can retain more information of cluster extent by highlighting ROIs with which each has notable overlap in a reference atlas. Table 2 shows this form of reporting for the same clusters in Table 1, which were calculated with AFNI’s “whereami” program and using the Glasser atlas as reference. Table 2 shows that 8 out of 11 of the primary clusters have >10% overlap with multiple atlas ROIs; this information would be lost in the standard single-voxel-summary approach. In some cases, the cluster peak and/or center of mass voxel are not even contained within a strongly overlapping ROI (e.g., Cluster #2, #3, #8 and #9). This highlighting approach retains much more useful information for each cluster when presenting results.

**Table 2.**
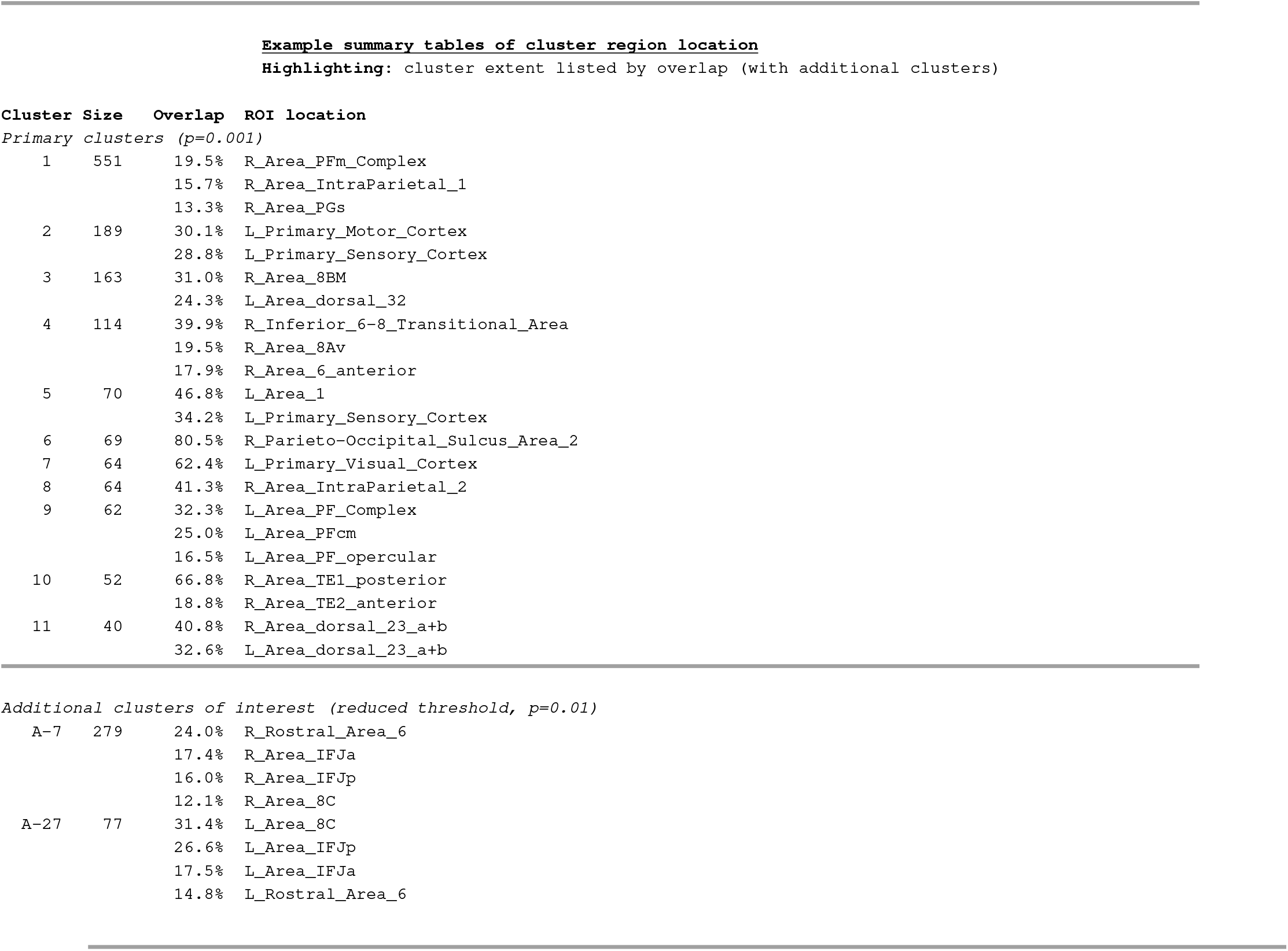
An example of summarizing voxelwise group results after clustering using a “highlighting” approach (with the Glasser atlas for reference location; cluster size refers to the number of voxels). Here, the set of atlas ROIs which comprise >10% of the cluster volume are shown (and this fraction could be adjusted). The main section of the table (top) shows the summary information for the suprathreshold clusters in Fig. 2A-C; the secondary section (bottom) shows how additional locations of interest that are visible with the highlighting approach can be presented (here, the two clusters with yellow arrows in Fig. 2, as determined by thresholding in 2D). Note how most clusters overlap meaningfully with multiple regions, each of which are highlighted by relative fraction. This can be contrasted with using a single voxel to summarize a voxel, as in Table 1. We see by comparison that neither the center of mass voxels for Clusters #2, #3 and #9 nor the peak voxels for Cluster #8 lie within a major region of overlap.

Additionally, we show an example of providing an extension of the cluster summary table, where additional clusters that were observed with the transparent thresholding can be listed. For example, these might coincide with the researchers’ pre-analysis hypotheses or with known physiological regions of interest, and hence be appropriate to comment on. Here, the two additional regions highlighted in Table 2 are symmetric (see yellow arrows in Fig. 2) and each overlaps strongly with the inferior frontal junction (IFJ) and areas in the precentral sulcus (Amunts et al., 2010). These areas have been strongly associated with “task difficulty” and “interference” control in neuroscientific literature (using Neurosynth, Yarkoni et al., 2011; and see Brass and von Cramon, 2004), as well as gambling (Li et al., 2010); as such, they may be appropriate to discuss further. This approach is in line with the highlighting strategy for more full results reporting (and the subthreshold nature of the additional regions is still clear).

Finally, we note that even though the atlas overlap is a much more informative approach of cluster localizing than using a single voxel, and therefore Table 2 is a preferable style of results reporting than Table 1, one should still avoid utilizing the table summaries when performing meta-analyses. Cluster tables have a dependence on threshold value(s), on the chosen minimal fraction of ROI overlap and on the choice of reference atlas (of which there are typically many to choose from). As such, they should only be used for providing approximate location/overlap information for reference within a study. Instead, cross-study comparisons and meta-analyses should be based on the unthresholded effect and statistical datasets, since those won’t be biased by arbitrary threshold values.

### Regionwise (ROI-based) results

For the region-wise analyses, a basic 2-sided t-test was performed on each of the 87 Desikan-Killiany regions, in a massively univariate approach. This produced mean and standard error estimates for the effect in each region, and these are represented using box-and-whisker plots in Fig. 4A. For each region, the color represents the relative amount of the effect estimate distribution that is positive, as reflected by the statistical evidence *E*_s_: therefore, stronger evidence for positive (or negative) effects are shown as more magenta (or green), with weak evidence being gray. The t-test coloration reflects FDR adjustment of statistical significance.

Also shown in Fig. 4A are box-and-whisker plots derived from the RBA-estimated posterior distributions. Comparing the results for each ROI, one observes that many median values and distributions are similar, with those from the RBA approach being typically (but not universally) smaller---this is expected from the data pooling aspect of the hierarchical modeling approach. However, the RBA results from the single model do not require a post-hoc multiple comparisons adjustment, which can be performed in different ways and is sensitive to a number of factors. With RBA, the ROI-wise results can be interpreted and presented directly without the need for any multiplicity correction.

We note that an important aspect of complete results reporting is model validation. This is particularly important when multiple methods can be applied, to be able to understand the relative benefits of one or the other; one cannot just choose the method with higher statistics or larger numbers of significant results, for example. Here, posterior predictive checking (PPC) and cross validation can be used to verify model quality and to compare model performance using Monte Carlo simulations. Fig. 4B shows the PPC fits for both the t-test and the hierarchical RBA modeling.^6^ By comparing the simulation curves (shown in amber) with the data curve (shown in black), it is apparent that the hierarchical approach provides a much higher model quality. In addition, one can more accurately compare the two models through quantitative indices such as various information criteria. For example, one can put each model through leave-one-out cross-validation and compare the expected log predictive density (ELPD) of each (Vehtari et al., 2017). As a rule of thumb, a model is preferable if its point estimate of higher ELPD is larger than two standard errors. Compared to the t-test, the hierarchical model showed a much higher predictive accuracy based on the expected log predictive density, with a difference of 1330 ± 50, suggesting the latter is a strongly preferable model.

These ROI results can be viewed as brain images in multiple ways, as shown in Fig. 5. The t-test (FDR adjusted) and RBA results are shown in panel A with opaque thresholding (here, at |*E*_s_| > 0.95). Even though FDR correction is not the strictest statistical adjustment approach, the RBA output shows more regions with strong statistical evidence at this level. While the supra-threshold regions are similar to those voxelwise results in Fig. 2A, but the opaque thresholding makes a reliable and informative comparison difficult, beyond perhaps counting the number of regions appearing (3 from the t-test, 6 from RBA).

Fig. 5B shows the same results with the highlighting approach, using transparent thresholding. The results from the two different statistical tests show a great deal of similarity with each other, and can more easily be compared with Fig. 2B, to which they also show a great deal of similarity. One can observe how much information is lost in the opaque thresholding. Summarizing the results with either just the few ROIs from the opaquely thresholded image, which would be the traditional practice, seems like an unnecessarily limited representation of the full parcellation analysis considering Fig. 5B (as well as Fig. 4A) and might well lead to different conclusions.

One notable difference in the ROI-based results from the earlier voxelwise ones is that the extreme effect estimates are generally lower in the ROIs, which is reflected in the difference in the range of the colorbars in Fig. 5 and Fig. 2. This is primarily due to the mixing of heterogeneous effects through averaging within the regions, particularly within the larger ones.

One way to decrease within-ROI heterogeneity is to use smaller regions. Here, we performed a second region-based analysis using the Glasser atlas (Glasser et al., 2016), which has 360 ROIs across a similar volume as the Desikan-Killiany atlas, which has 87 regions. We applied the same t-test and RBA approaches, the results of which are shown in Fig. 6. More details are visible within the cortex, and the same relative properties between the t-test and RBA results are observable. With this a finer parcelation, the effect estimates are closer to those of the voxelwise testing (though still noticeably smaller).

**Figure 6.**
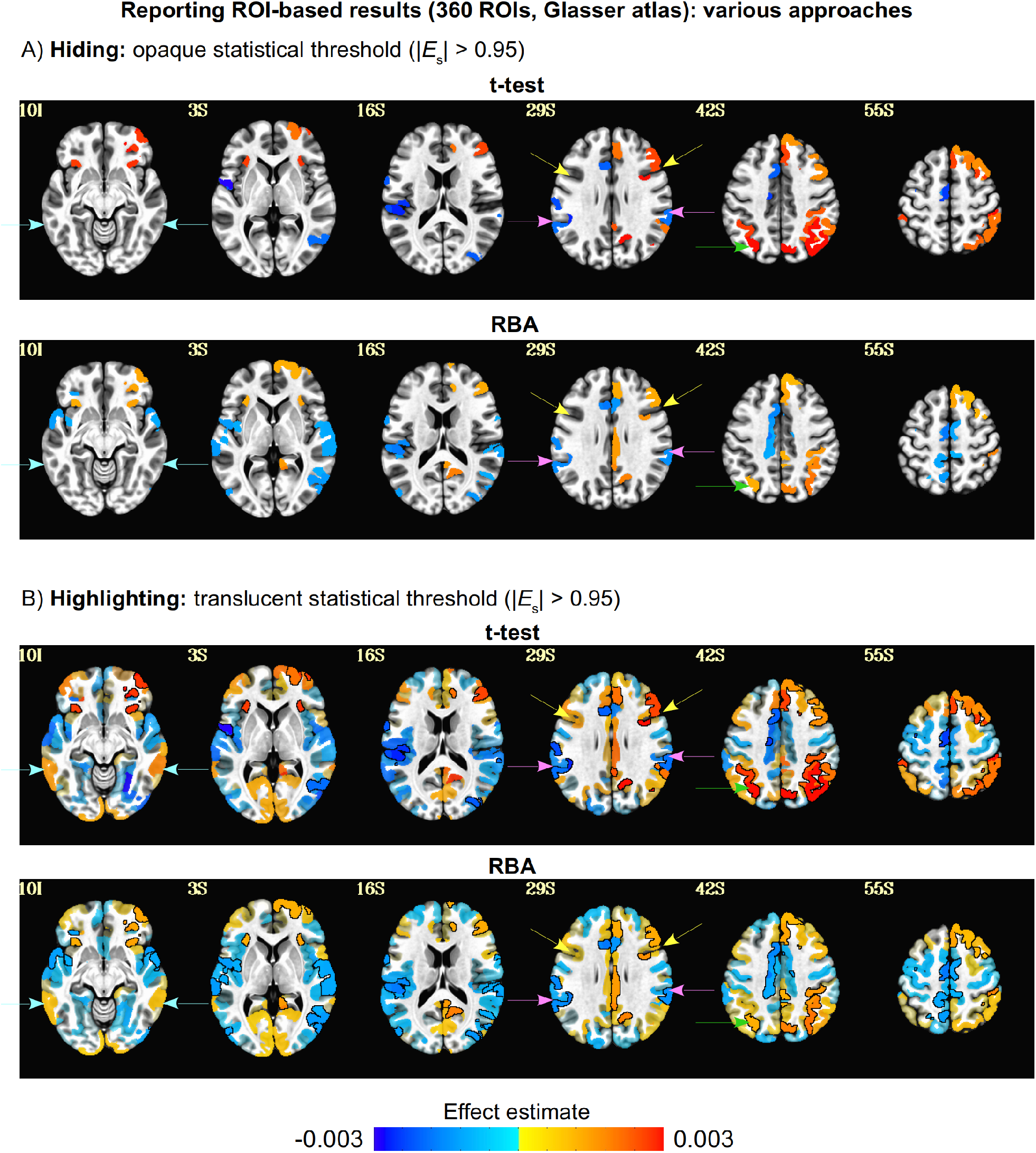
A comparison of group-level effect estimation for ROI-based results using the traditional style of “hiding” results and the newer “highlighting” style (axial slices: left=left). The modeling and presentation style for each panel are the same as used in Fig. 5, but here using the 360 ROI Glasser atlas (Glasser et al., 2016). The results are generally similar to those using the Desikan atlas, but the range of effect estimates tends to be wider for both the t-test and RBA results. The extreme effect estimates here are still notably smaller than the voxelwise case, but they are closer than the results of the Desikan atlas, likely due to having smaller ROIs and less heterogeneity per ROI.

We note that when presenting all of the above regionwise and voxelwise outputs, we have shown the estimated effects (or “point estimates”) as overlays, rather than the test statistics. The latter is just statistical information that is already used for thresholding. While the same brain locations would be shown in each case, coloring the voxels by effect---rather than redundant statistical information---provides complementary information in the reporting. In most areas of science, the focus is on the effect itself, and not on the statistic, since the former should have physical meaning and the latter is a dimensionless value for statistical evidence. Moreover, the estimated effect provides a vehicle for more accurate meta-analysis.

### Cross-study (“cross-team”) results

Finally, we compare all-or-nothing reporting with the “highlighting” approach when assessing the consistency of results across several studies. Fig. 7A shows the set of 61 statistics maps^7^ that apply to both Hyp #1 and 3 in the NARPS project (the “equal indifference” group, for the gain effect), using standard opaque thresholding. The threshold was set at |Z|,|t|=3 (again, the NARPS teams’ results could be reported in either statistic, but these should be approximately equivalent, given the *N ≈* 50 subjects in the group results). In Panel A, the largest pattern of similarity is the large negative region in the superior frontal cortex (visible in many brains in the first four rows), which appears in perhaps half of the results, though with varied shape and extent. Cluster overlap appears to be high for a few datasets, but in general the variability of cluster shape, extent and existence appears to be relatively high. This can be seen quantitatively in Panel B, which shows similarity matrices among all the brains using the Dice coefficient of similarity (appropriate here given the dichotomization of opaque thresholding). These hypotheses focused specifically on the positive-effect clusters, but we also show the similarity matrices for both positive and negative clusters to provide a complete picture.

**Figure 7.**
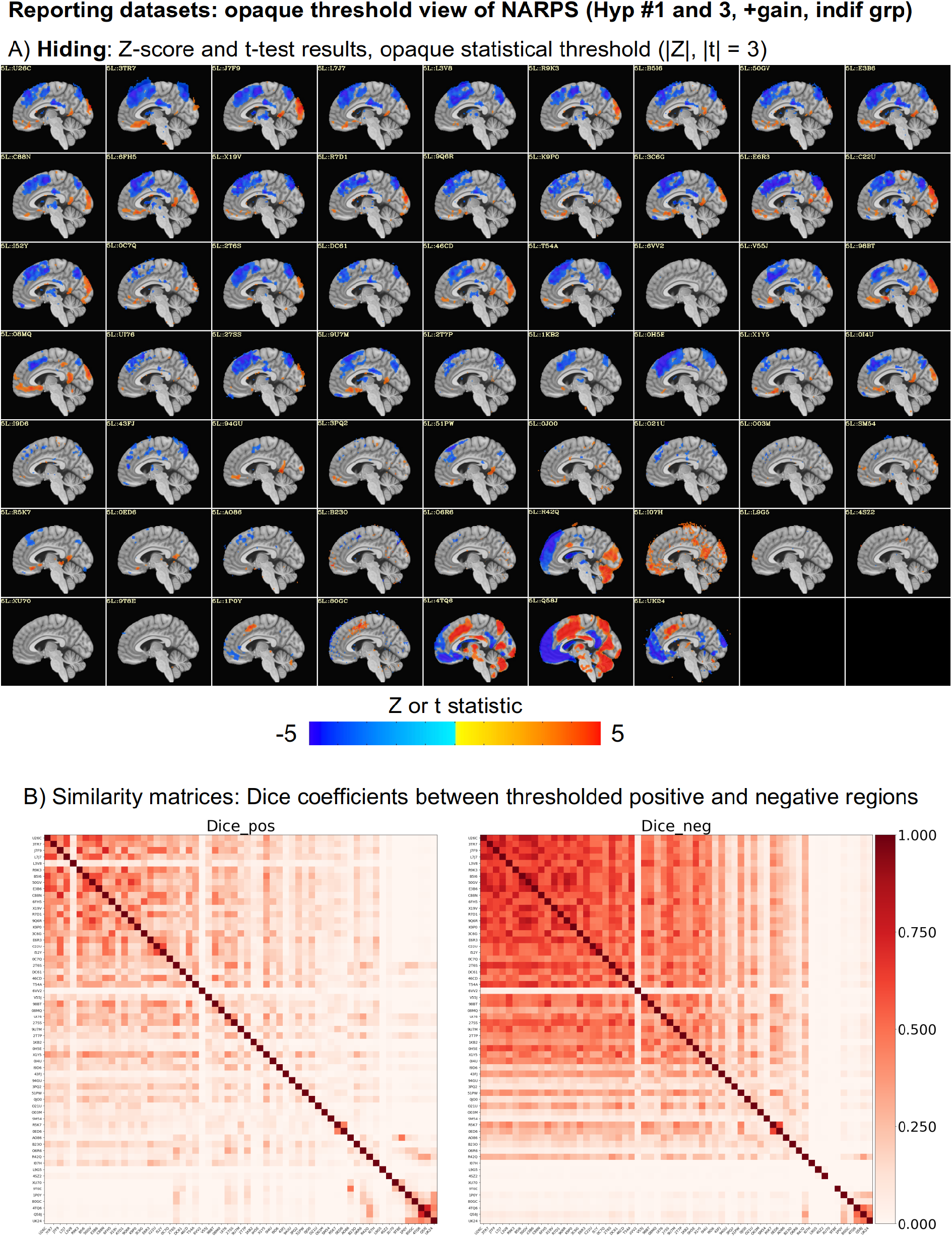
Cross-study meta-analyses using the traditional style of “hiding” results “(sagittal slices: left=anterior). Panel A shows thresholded maps of submitted results from NARPS research participants for Hyp #1 and 3 (“equal indifference” group, for positive “gain” effect, which had lowest agreement in the dichotomized part of the NARPS reporting), downloaded from NeuroVault: each slice contains voxelwise t- or z-statistic results (which are essentially equivalent, given the number of approx. N=50 subjects in the group) thresholded at |t|,|z| = 3. Panel A shows a great deal of heterogeneity, since the thresholded islands obviously differ across a large number of results. This is shown using similarity matrices of Dice coefficients for each of the set of positive and negative regions in Panel B. See Fig. 8 for the same results presented in “highlighting” mode.

**Figure 8.**
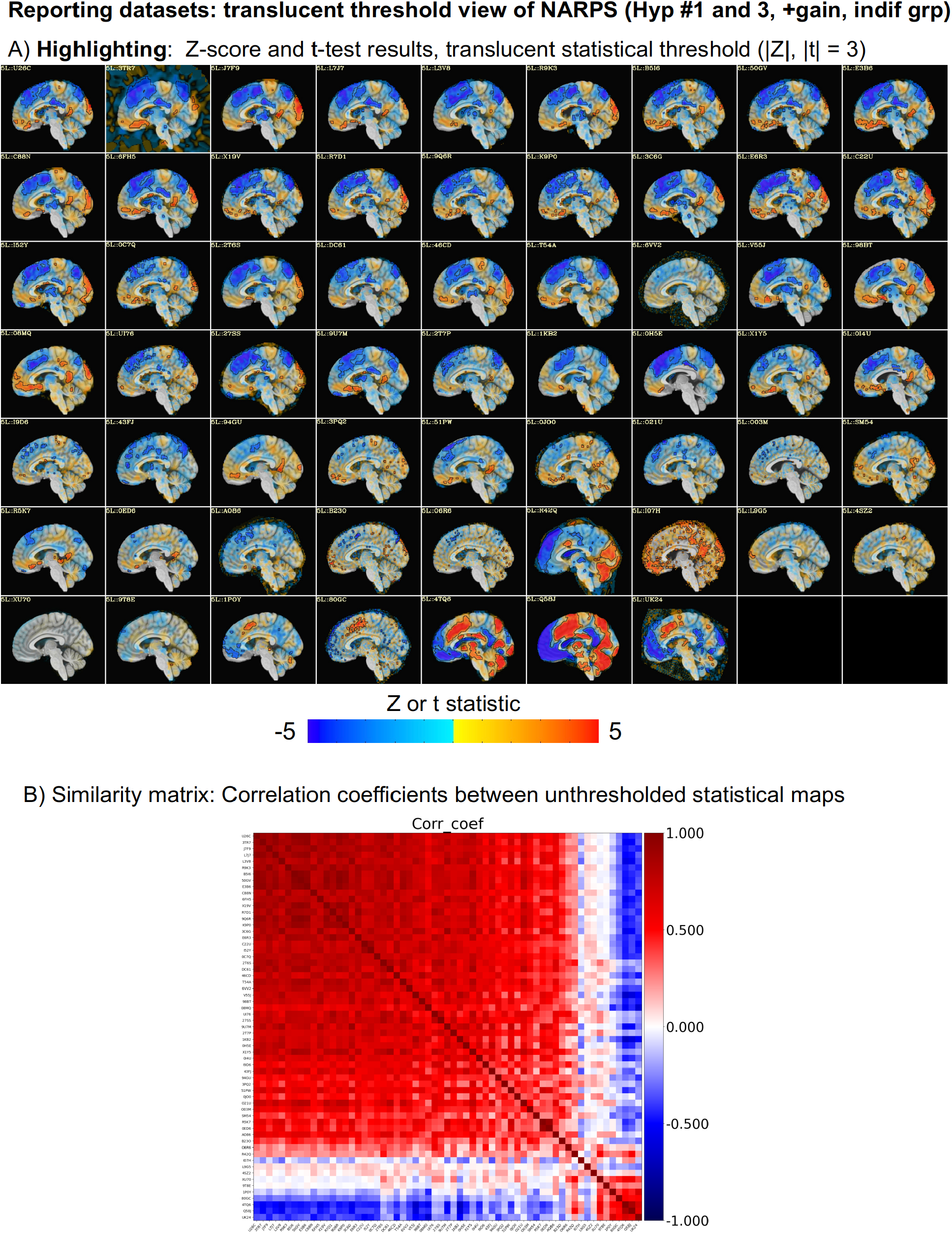
A presentation of the same results shown in Fig. 7 but using the “highlighting” style. In contrast to the opaquely thresholded results, here Panel A shows that the vast majority patterns of positive and negative effects are actually quite similar across groups, while just the magnitudes of the statistics differ (though, indeed a very small number of brains differ in notable ways; see text for details). The transparent thresholding provides a more accurate assessment of results, and points to a specific mode of variation (e.g., strength of results from some participants) rather than blanket statement of “wide variation”, which is not accurate, from dichotomized results. This is shown by the corresponding similarity matrix using Pearson correlation coefficients across the whole brain maps (consistent with the “highlighting” approach of using more data than thresholded values).

Fig. 8A shows the same set of statistical maps using the transparent thresholding approach. This style of reporting shows much more information for each team’s result, and it suggests a very different level of cross-study similarity: the patterns of large positive and negative statistics appear to be quite similar across a large fraction of the brains (approx. 51 “Block 1” datasets, out of 61 total), compared with the binarized maps shown in Fig. 7A. While the statistical strength varies and may be sub-threshold in varying degrees, this display with more information content shows a very large degree of general agreement in both locations and directionality of results. That is, what varies in these images is primarily the scale of the statistical values, rather than the signs or patterns. This suggests that a major driver of the cross-teams differences is the statistical power of their methods, rather than differences in the actual effect sizes they are estimating.

The corresponding similarity matrix between the unthresholded maps can be calculated with Pearson correlation between the dataset pairs, shown in Fig. 8B. This reinforces the high degree of similarity across most teams’ results, in contrast to those of the hiding approach shown in Fig. F7B.

Additionally, one can observe a second, smaller block of high inter-team agreement among the final brains that is also strongly anticorrelated to the main block (approx. 5 “Block 2” datasets); this is denoted by the dark blue rectangular regions, and also verified in the images in panel A. Thus, this block appears to differ from the main one primarily by a sign flip in the reported results, either by an intentional sign convention or by mistake in the processing and analysis. This could be caused by taking the incorrect file in a pair of 1-sided testing results, for example (an error that can be easily avoided by implementing 2-sided testing). In fact, the same block of approx. seven team’s results were observed and commented upon in the original NARPS paper, who attributed it to: misspecification of modeling in four cases; the inclusion of multiple regressors correlated with the gain parameter in two cases; and unknown reasons in one. These differences in results are important to understand, and we emphasize that viewing the unthresholded results is what allowed them to be noticed both here and in the original NARPS paper.^8^

Excluding this sign-flipping issue, only a very small fraction of the teams’ results have low correlation to these two blocks (the 5 remaining “Non-block” datasets). The vast majority of results (56/61 = 92%) agree strongly when opaque thresholding is not imposed. In the thresholded results, the fraction of agreement would be viewed much lower: the blocks of high-Dice values in matrix Fig. 7B are much smaller, particularly for positive-clusters. Note that there is no processing difference between the results in Fig. 7 and 8, only the choice of whether to add a thresholding step before making the comparison. The information lost from opaque thresholding greatly reduces the apparent similarity, while transparent thresholding does not do so.

We note that choice of software package did not appear strongly associated with being in a particular block. Table 3 shows the following information for each dataset in Figs. 7-8 (in the same order): software package used; whether fMRIPrep (Esteban et al., 2019) specifically was used in the early stages of processing; test approach; and an estimation of smoothness of Hyp. #1 statistics volumes. The horizontal lines in the Table delineate blocks. Note that combinations of software choices appear scattered throughout the table, and each block contains a mix of packages and other processing choices. For example, Yes/No answers for fMRIPrep usage and Par/NonPar answers for testing methods appear in each of Block 1, Non-Block and Block 2.

**Table 3.** The list of participating teams’ softwares and testing used (as reported in Ext Table 2 of Botvinick-Rezer (2020)), in the same order as the figures and correlation matrix rows/columns in Figs. 7-9. The horizontal lines denote the breaks between the 51 “Block 1”, 5 “Non-block” and 5 “Block 2” datasets, respectively. Note that combinations of software choices appear scattered throughout the table, and each block contains a mix of packages and other processing choices. For example, Yes/No answers for fMRIPrep usage and Par/NonPar answers for testing methods appear in each of Block 1, Non-Block and Block 2. Idx = index in order of results; Team = NARPS team ID; Pkg = software package used; fprep? = was fMRIPrep included in analysis; Test = was parametric (Par) or nonparametric (NonPar) used; SmooH1 = a smoothing value estimated for statistical results of Hyp. #1 (and therefore also for Hyp. #3) with FSL’s “smoothest” program.

| Idx | Team | Pkg | fprep? | Test | SmooH1 | Idx | Team | Pkg | fprep? | Test | SmooH1 |
| --- | --- | --- | --- | --- | --- | --- | --- | --- | --- | --- | --- |
| 1 | U26C | SPM | Yes | Par | 10.38 | 37 | I9D6 | AFNI | No | Par | 6.21 |
| 2 | 3TR7 | SPM | Yes | Par | 17.40 | 38 | 43FJ | FSL | No | Par | 10.66 |
| 3 | J7F9 | SPM | Yes | Par | 14.88 | 39 | 94GU | SPM | No | Par | 11.19 |
| 4 | L7J7 | SPM | Yes | Par | 11.76 | 40 | 3PQ2 | FSL | No | Par | 5.79 |
| 5 | L3V8 | SPM | No | Par | 14.74 | 41 | 51PW | FSL | Yes | Par | 11.15 |
| 6 | R9K3 | SPM | Yes | Par | 11.77 | 42 | 0JO0 | Other | Yes | Par | 8.12 |
| 7 | B5I6 | FSL | Yes | NonPar | 9.84 | 43 | O21U | FSL | Yes | Par | 8.26 |
| 8 | 50GV | FSL | Yes | Par | 10.26 | 44 | O03M | AFNI | Yes | NonPar | 3.47 |
| 9 | E3B6 | SPM | Yes | Par | 12.80 | 45 | SM54 | Other | Yes | Par | 7.05 |
| 10 | C88N | SPM | Yes | Par | 11.62 | 46 | R5K7 | SPM | No | Par | 12.06 |
| 11 | 6FH5 | SPM | No | Par | 12.22 | 47 | 0ED6 | SPM | No | Par | 7.86 |
| 12 | X19V | FSL | Yes | Par | 8.48 | 48 | AO86 | Other | Yes | NonPar | 7.49 |
| 13 | R7D1 | Other | Yes | NonPar | 8.93 | 49 | B23O | FSL | Yes | NonPar | 3.32 |
| 14 | 9Q6R | FSL | No | Par | 10.28 | 50 | O6R6 | FSL | Yes | NonPar | 3.06 |
| 15 | K9P0 | AFNI | Yes | Par | 8.05 | 51 | R42Q | Other | No | Par | 12.73 |
| 16 | 3C6G | SPM | No | Par | 14.26 | 52 | I07H | Other | Yes | NonPar | 5.59 |
| 17 | E6R3 | Other | Yes | Other | 9.28 | 53 | L9G5 | FSL | No | Par | 7.22 |
| 18 | C22U | FSL | No | Par | 11.16 | 54 | 4SZ2 | FSL | Yes | Par | 6.65 |
| 19 | I52Y | FSL | No | NonPar | 11.42 | 55 | XU70 | FSL | No | Par | 7.17 |
| 20 | 0C7Q | Other | Yes | NonPar | 8.68 | 56 | 9T8E | SPM | Yes | NonPar | 9.85 |
| 21 | 2T6S | SPM | Yes | Par | 14.93 | 57 | 1P0Y | SPM | No | Par | 9.13 |
| 22 | DC61 | SPM | Yes | Par | 9.58 | 58 | 80GC | AFNI | Yes | Par | 4.02 |
| 23 | 46CD | Other | No | Par | 10.92 | 59 | 4TQ6 | FSL | Yes | NonPar | 14.88 |
| 24 | T54A | FSL | Yes | NonPar | 12.28 | 60 | Q58J | FSL | No | Par | 16.24 |
| 25 | 6VV2 | AFNI | No | Par | 7.20 | 61 | UK24 | SPM | No | Par | 10.76 |
| 26 | V55J | SPM | No | Par | 12.85 |  |  |  |  |  |  |
| 27 | 98BT | SPM | No | Par | 11.48 |  |  |  |  |  |  |
| 28 | 08MQ | FSL | No | NonPar | 13.14 |  |  |  |  |  |  |
| 29 | UI76 | AFNI | Yes | Par | 6.60 |  |  |  |  |  |  |
| 30 | 27SS | AFNI | No | Par | 11.37 |  |  |  |  |  |  |
| 31 | 9U7M | Other | No | Par | 14.78 |  |  |  |  |  |  |
| 32 | 2T7P | Other | No | Other | 7.66 |  |  |  |  |  |  |
| 33 | 1KB2 | FSL | No | Par | 13.06 |  |  |  |  |  |  |
| 34 | 0H5E | SPM | No | Par | 14.17 |  |  |  |  |  |  |
| 35 | X1Y5 | Other | Yes | NonPar | 8.69 |  |  |  |  |  |  |
| 36 | 0I4U | SPM | No | Par | 8.69 |  |  |  |  |  |  |

Since NARPS was specifically focused on particular regions of interest, we investigated the cross-team similarity matrices for a subregion of the brain that encompassed the VMPFC and VST (the locations of interest for Hyp #1 and #3, respectively). These matrices are shown in Fig. 9. The Dice coefficients of the hiding approach in matrices A-C show even lower similarity than the whole brain results in Fig. 7B. In contrast, the correlation coefficients of the highlighting approach in matrix D shows even higher similarity than the whole brain results in Fig. 8B (again, accounting for sign flips).

**Figure 9.**
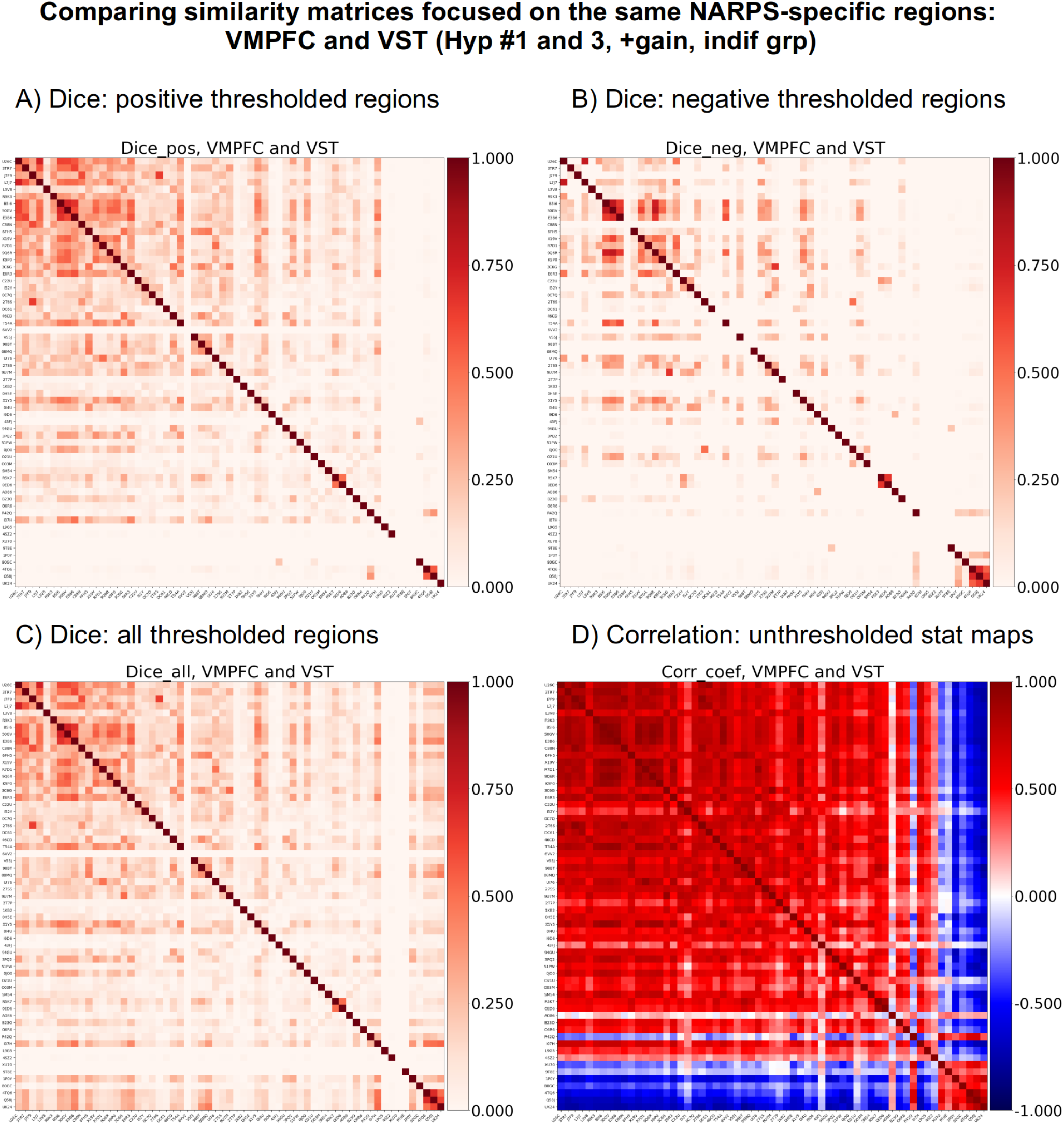
A presentation of the same results shown in Figs. 7-8 for a zoomed-in sub-region enclosing the ROIs of interest for Hyp #1 and #3 (which share the same underlying data). Matrices A-C reflect the “hiding” approach of comparing results across the group, while matrix D reflects the “highlighting” approach. Similar to the whole brain comparisons in Figs. 7-8, there is a notable difference between the highlighting and hiding approaches. Here, in the locations of most interest for this particular data, the difference is even more apparent: the correlation approach, which retains much more information from the analysis, shows a great deal of similarity across the teams’ results. Even the secondary cluster of similar results is shown to be strongly related to the results of the initial processing, differing mainly in sign (e.g., due to a coding error, an analysis choice, etc.). Only one or two results appear to be unrelated to the main clusters.

An interpretation of results based on Fig. 7’s dichotomized presentation might emphasize high variability (particularly if the results were further thresholded to be 1-sided, rather 2-sided), while one based on Fig 8’s more complete results reporting might emphasize similarity. Note that the latter approach includes a higher degree of information, and that this occurred at both the whole brain scale and when focused on the hypotheses’ regions of interest. These potentially contradictory conclusions demonstrate the importance of preserving data integrity as much as possible in result reporting, in order to provide accurate and complete assessments.

## Discussion

### The problems with traditional “hiding” of results, and the benefits of the “highlighting” approach

There are many problems that arise from standard results reporting methodology that hides so much of the data. We list these here, and show how each can be addressed and ameliorated by the highlighting framework.

#### 1) Unrealistic ON/OFF representation of biology

Traditional results reporting promotes interpreting “brain activation” as binarized: that is, reporting that some contrast or activity was found in (a small number of) clearly delimited regions X and Y, and absolutely nowhere else. However, this is almost certainly not a physiologically-realistic interpretation of FMRI measures of brain behavior. For example, Gonzalez-Castillo et al. (2012 and 2015) scanned subjects with a large number of runs (100 each) and repeatedly showed that the extent of activation areas monotonically increased as data from more runs were included in the analysis. Traditional thresholding cannot capture this reality of the BOLD response, while the highlighting approach can. This latter approach is also more in-line with the increasing number of neuroscience approaches that focus less on individual regions in isolation and more on interactions and the full context of brain behavior and activity.

#### 2) Unrealistic null hypothesis significance testing interpretation

Perhaps surprisingly, opaque thresholding could be viewed as violating the basic rules of the standard, widely used NHST with which it is so closely associated. Briefly, one establishes some null hypothesis H_0_ (e.g., the mean effect μ = 0) and an alternative hypothesis H_a_ (e.g., μ ≠ 0). Then, after modeling one decides whether to reject H_0_ in favor of H_a_ at a predefined significance level. The NHST includes two important rules of interpretation: 1) rejecting H_0_ does *not* prove that H_a_ is true, and 2) *not*-rejecting H_0_ does *not* prove that H_0_ is true. However, the standard practice of zeroing out sub-threshold regions (places where H_0_ is *not* rejected) is effectively proclaiming H_0_ to be true there, because any contrary evidence is hidden. This violates Rule #2. Are we really 100% certain that there is *exactly no effect* there---and therefore those results can be zeroed---even though the NHST itself cannot support that interpretation? It would be more statistically appropriate to present the evidence that exists there.

#### 3) Within-study information waste

Related to the prior point, suppose a null hypothesis of activation in a particular region X had been made, but then a cluster was not found there to reject it. With opaque thresholding, one has no idea *why* a location failed to reach the chosen threshold in that study--was the response in that area near or far from the designated evidence strength cut-off, and in what direction? Was there a sizable effect magnitude that was washed out by very large variability, or an effect just a bit too small and larger sample sizes (or a more appropriate model) might provide more conclusive evidence? Recall the important axiom: *the absence of (strong) evidence is not the same as the (strong) evidence of absence*. This is all important information for truly understanding a study’s outcomes, but it becomes wasted by standard, thresholded visualizations (Chen et al., 2022b). We should not use figures to pretend that there was “no evidence” when in fact some was present, and the highlighting approach retains it.

#### 4) Unrealistic view of noise

FMRI data are noisy, but the noise is not homogeneous. Within a single scan, noise will vary spatially based on MRI receiver coil locations, distortions, and drop out. Noise from motion, cardiac and respiratory effects are also not uniform. Across participants, there is additional noise from slice coverage and spatial registrations. These ubiquitous characteristics of FMRI data greatly affect the extent and location of high-signal and high-significance regions: for a given study, certain brain areas will reliably have more noise than others. In practice, this means that while the superior temporal sulcus (STS) and the temporal pole might each show a 1% effect size in a study, the former might pass thresholding and the latter, not. Knowing that the temporal pole signal is noisier, it would be inappropriate both to overstate the importance of the STS finding and to totally ignore the temporal pole observation. Most experiments nowadays probe subtle effects (≲0.1-1%) with event-related designs, as well as the contrasts between these signals. Those are small effects in a sea of noisy data that contains spatially inhomogeneous distortions. As opaque thresholding is highly sensitive to threshold values and local noise patterns, it might easily hide real effects or lead to overstating the importance of a few regions that cross threshold when the full context is not shown (by ignoring other/secondary regions). The highlighting approach avoids these issues, benefitting scientific interpretation.

#### 5) Interesting features can be hidden

Opaque thresholding can also hide meaningful patterns and features within the data. Allen et al. (2012) presented a case study analyzing an event-related FMRI paradigm with an auditory oddball task. With a standard visualization approach that used opaque thresholding of a contrast for a 1-sided hypothesis, expected regions were observed as statistically significant (e.g., auditory cortex). However, when viewing results with transparent thresholding (and looking at 2-sided test results; see Chen et al., 2019a), the authors clearly observed the additional feature of a default mode network pattern. While FMRI studies focus on predetermined hypotheses, our knowledge of brain dynamics is not perfect, and the highlighting visualization allows for observation of unexpected results (which, of course, require further testing for reproducibility, etc. before being fully accepted) that may help formulate novel hypotheses.

#### 6) Biases in thresholding

Small anatomical regions are consistently discriminated against by standard cluster-based approaches due to their intrinsic size disadvantage, even if their effect magnitude or statistical evidence is comparable or even greater than that of larger clusters. The survival difficulty of such small regions through whole-brain analysis may force the investigator to seek for band-aid solutions such as reducing the data domain to a list of predefined regions and small volume correction for the sake of passing the designated threshold, potentially wasting the data information in the rest of the brain. While some FMRI group analysis techniques have been developed specifically to reduce dependence on individual voxelwise thresholds, such as TFCE (Smith and Nichols, 2009) and ETAC (Cox, 2019), there is still notable threshold dependence in intermediate steps. The highlighting methodology reduces the need for alternative strategies that might incur other trade-offs, or at least it could be applied to these outputs, as well.

#### 7) Equal threshold values do not ensure equal evidence

Consider an ideal meta-analysis of typical studies reporting on the same set of brain regions, each using the same “standard” thresholds. Even in such a scenario, each study will generally *not* be providing equally weighted evidence. For example, subject sample size will vary (potentially by orders of magnitude, in modern neuroimaging), as will SNR and noise/distortions since the data were collected on different scanners and potentially at field strengths ranging from 1.5-7T, with different parameters, shimming, coils, etc. The FMRI data were also likely acquired under different conditions: task studies could be a mixture of block design, event-related and amplitude-modulated paradigms, each using different stimuli and various trial sample sizes; naturalistic designs could use different movies or songs played to subjects; resting state studies have varied duration and TR. Those create inherent differences among the data collections themselves.^9^ All-or-nothing thresholding will be sensitive to those aspects and result in biased views of the results; highlighting would provide a much fairer way of evaluating across sites.

#### 8) Incorrect effect magnitudes and signs

Mathematically, only reporting statistically significant results may bias the effect magnitude toward overestimated sizes and/or incorrect directionality (Gelman and Carlin, 2014). In other words, a stringent focus on false positive rates can lead to reporting effects with inflated magnitudes (“Type M” error) and/or incorrect signs (“Type S” error). This feature is related to the classic “publication bias” problem, wherein findings with weak or negative evidence do not get reported, which biases meta-analyses (Sterling, 1959; Ioannidis, 2008; Jennings and Van Horn, 2012). Furthermore, it connects with the importance of reporting effect estimates in neuroimaging studies (Chen et al., 2017): misestimation of effect magnitudes and signs can easily lead to misinterpretation and to mistakes in replication. Transparent thresholding reduces this bias, and increases the chances that misestimation will be observed and addressed.

#### 9) Arbitrariness features prominently in results

In theory, NHST thresholding is a rigorous, quantitative procedure, so dichotomization is appealing for its objectivity. However, in practice it contains several elements of arbitrariness. One cluster may fail at an FWE of 0.05, but survive at FWE of 0.055--why is the former an inherently “better” filter of outputs (just because it is a rounder number in base 10)? A region may just fail FWE correction with one acceptable modeling method, but just pass with an equally acceptable one such as FDR or Bayesian multilevel modeling---which very different result is correct? Two clusters may have the same effect size, but due to noise/uncertainty one is slightly disjoint (and fails thresholding)---should we see nothing of that evidence? In each case small contingencies can dramatically affect the dichotomized results and negatively impact the study’s reliability. Some have argued that these issues could be addressed by picking stricter limits for threshold values, such as moving the “standard” threshold from FWE=0.05 to FWE=0.005 (Benjamin et al., 2018). However, all the same sensitivity issues will still apply, just for FWE=0.0055 vs 0.0049, etc. Moreover, it has been shown that there is no simple or direct connection between a designated significance threshold and the likelihood of similar outcomes in subsequent studies (McShane et al., 2019). Consider the proposed alternative: transparent thresholding would display the similar results with appropriately small differentiation. This was demonstrated here, with the apparent differences between the t-test- and RBA-based analyses were greatly reduced when presented with highlighting; even the differences between voxelwise- and region-based analyses were shown to be largely in agreement when highlighting.

#### 10) Reproducibility issues

Consider that region X has been the focus of ten studies. Positive results crossed standard significance thresholding in only 2 of them, but there were also moderately significant, sub-threshold positive results observed there in 5 others. Under current practices, the latter evidence would be entirely absent from all reporting and a meta-analysis would list 2 of studies “for” positive activation there and 8 “against.” The activation effect would be labeled as irreproducible. However, that is likely not an accurate assessment of the results of all those studies. Moreover, what if those five sub-threshold results showed the *opposite* effect there? That important counter-evidence, which would be useful for accurate interpretation, would also be hidden. Dichotomization of results may be “clean”, but it wastes important evidence across studies and potentially leads to false conclusions, such as “nothing happening here”, instead of accurately assessing the additional evidence for/against (“things often here, with varied strength”). Transparent thresholding and reporting would lead to more accurate estimations of reproducibility---that is, one can observe where studies really do tend to conflict or where they provide similar evidence but with varied degrees of weight. The comparison of NARPS teams’ results is a good example of how dichotomization can make relatively small variations in agreement look divergent, while transparent thresholding allows for a more accurate assessment.

#### 11) P-hacking

In theory, having a “standard” threshold should reduce or eliminate p-hacking. However, in practice, all-or-nothing dichotomization may actually encourage the use of other, non-threshold-based manipulations of statistical evidence, which are distinct from tests of analytic robustness. This can include implementing several statistical tests or testing various combinations of model factors until some (but not too many, or too big) clusters survive the standard threshold. While the p-hacking problem has been noted previously and some solutions put forth (e.g., Wicherts et al., 2016), including study preregistration (Nosek et al., 2018), the current practices of results reporting still provide a strong (and unfortunate) motivation for it. Transparent thresholding makes it possible to discuss potentially interesting sub-threshold observations and thus reduce the incentive for shuffling around statistical analyses (which has other possible ramifications) with the primary goal of trying to convert a slightly sub-threshold result into something significant.

#### 12) Artifacts can be hidden

Among the valuable pieces of information that may be hidden by opaque thresholding are quality control (QC) issues, either at the single subject or group level. Scanner artifacts or ghosting may be apparent in the patterns of non-statistically significant results. One may also observe processing-induced artifacts: ICA-based denoising might have mistakenly removed gray matter (GM) signals; masking may hide signs of ghosting; the shape of patterns may suggest poor alignment to a template space; etc. With opaque thresholding, one might not notice the presence of an underlying artifact, or that a peak of “significant” activation is actually due to an artifact whose tell-tale spatial pattern is hidden away^10^. With transparent thresholding, one is more likely to notice the presence of artifactual patterns, the presence of poor alignment, and more. See Box 1 for an illustrative example.

#### 13) Poor modeling can be hidden

Related to general QC issues is the question of model validation. Even at the best of times, modeling FMRI data is tricky, due to the large measurement noise scale, hierarchical levels of variability (e.g., cross trials, cross subjects) and other potential confounds such as scanner distortions and subject motion. To date, questions of optimal ways to denoise and model FMRI still remain, and both subject and trial sample sizes often remain relatively small (Chen et al., 2022a). Showing results from a wider set of regions/locations can give a better sense of the general appropriateness and quality of modeling. For example, seeing large patches of relatively high correlation following the edge of the brain may be a sign that subject motion effects had not been modeled well. Showing more data provides an indirect sense of model uncertainty and validation, which is important and useful in interpreting results. Furthermore, as demonstrated here, additional and more direct approaches for model validations exist and should be used in conjunction with this visualization (e.g., the PPC simulations from region-based modeling).

##### Box 1.

An example of the benefits of the transparent highlighting approach for quality control (QC) during processing. Brain-masking has also not been applied, for more full QC. The green arrow points to a sign of artifact that is hidden with opaque thresholding, but able to be observed with transparent highlighting.

###### Transparent thresholding to improve quality control (QC): a resting state FMRI example

When processing FMRI time series that do not have stimulus timing (e.g., resting state and naturalistic paradigms), AFNI’s afni_proc.py quality control (APQC) HTML report contains seedbased correlation maps of major networks. This helps both to check the data for artifacts and to verify various processing steps, such as motion estimation and regression modeling. The systematic images are created using transparent thresholding, to retain more information about the dataset and to reduce sensitivity to the threshold value, which has often proved useful.

Consider the two forms of the same correlation map in Panels A and B, below, which are from the same dataset, have the same threshold value of Pearson |r| > 0.3 and the same seed location in the posterior cingulate cortex. Panel A uses opaque thresholding, and one would likely interpret the results as a fairly standard picture of the default mode network; this subject would pass the QC check here. Panel B uses the transparent thresholding from the APQC report, and one quickly observes a large patch of negative correlation through frontal white matter (green arrow). While the anticorrelation strength is sub-threshold, the size, shape and location of the patch raise questions about the possibility of artifacts.

Indeed, further checks within this subject’s data (e.g., investigating correlation patterns with AFNI’s InstaCorr) reveal odd patterns of strong, spurious correlations. Panel C shows an example of this, with another correlation map from the same dataset using a seed located in the middle of the suspicious white matter region (center of the crosshairs). The strong correlation and anticorrelation patterns are likely artifactual and not physiological, and using the more informative transparent thresholding led to this discovery.

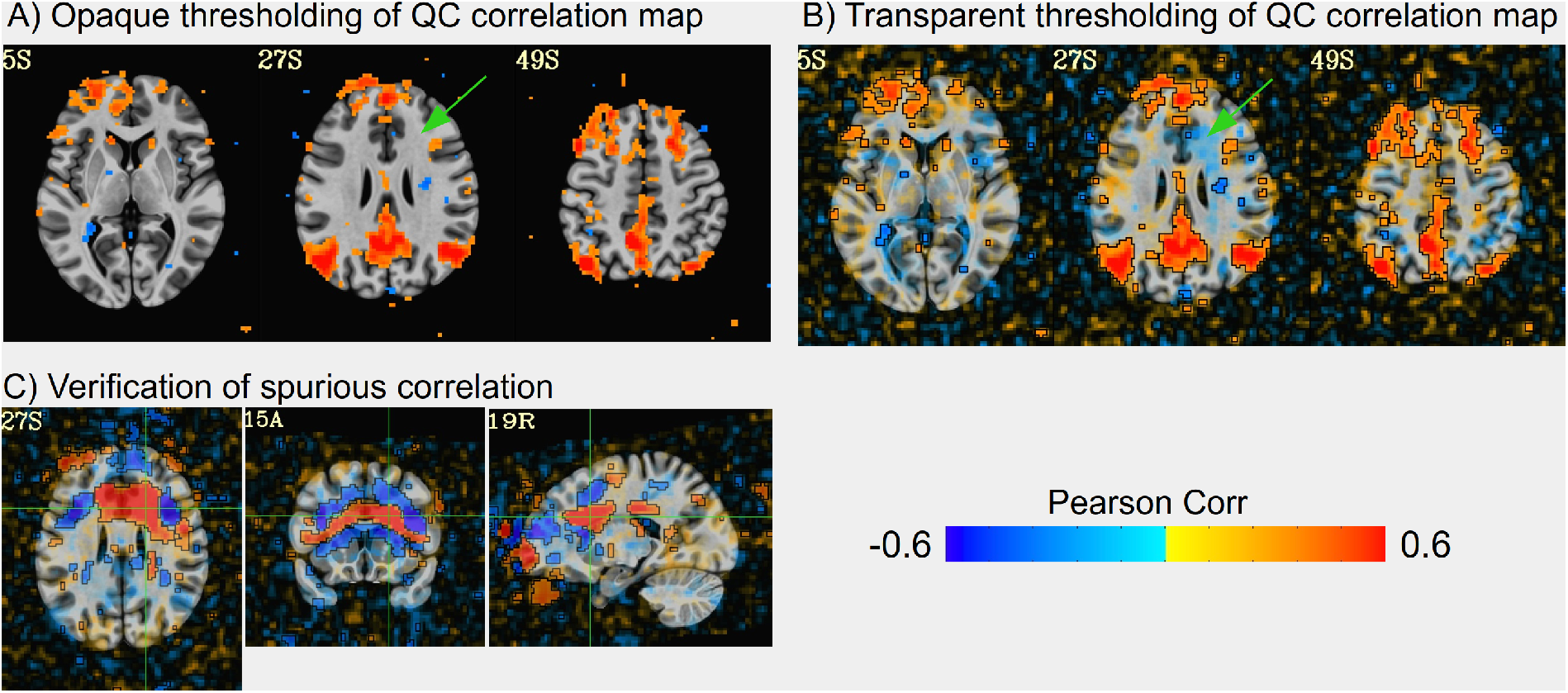

In summary, the originally desired properties of traditional results reporting with opaque, all-or-nothing thresholds---such as objectivity, reliability and reproducibility---are not met often in practice. Instead, the dichotomized and opaquely thresholded results form an artificial representation of activation, and one which is likely to have an inflated effect size. While thresholds are “objective” and “quantitative” in the sense of being based on a predefined number, in reality their values contain several subjective choices, which strongly affect results. There is no avoiding the arbitrariness of them. Most importantly, opaquely thresholded results do not contain all the information worth reporting in a study dataset. Potentially “real” effects may be hidden from the reader, and potentially artifactual ones may be unwittingly displayed. Worse yet, each of these scenarios may be hidden from the researchers themselves. Transparent thresholding in the “highlighting” approach alleviates these issues, benefitting scientific accuracy and interpretation.

### Recommendations for results reporting

To improve results reporting within the neuroimaging community, we would make the following suggestions when writing papers and making presentations. As a general guiding principle, individual studies should focus less on an “ON/OFF” mindset for reporting brain results, which opaque thresholding reinforces; instead, they should focus on presenting and providing evidence that can be combined with other studies. The benefits of many of these have been demonstrated here:

1. **Apply thresholds with transparency**. This “highlighting” mode reduces the sensitivity to arbitrary threshold values, increases the information available for interpretations, and leads to better representations of the data.
2. **Avoid summarizing voxelwise results as cluster peak locations**. That single value (or, equivalently, the center of mass) is generally a poor representation of complex data and biology, and it harms reproducibility efforts through selection biases and poorer meta-analyses.
3. **Avoid summarizing ROI-based results with only suprathreshold regions**. Provide tables that contain the evidence for all ROIs analyzed, including both effect estimation and uncertainties.
4. **Avoid masking results whenever possible**. Masking the data reduces the ability to observe potential artifacts or spurious features that affect results within the brain volume.
5. **Include and show more than just statistical results**. Effect estimates contain useful information for interpreting and comparing a study. Statistics are helpful for (transparent) thresholding, but they do not tell the whole story of the data. Wherever possible, provide model validations (e.g., as done in the regionwise analysis here and shown in Fig. 4B).
6. **Provide detailed meta-information about analysis**. This includes the (final) number of subjects, sidedness of testing, number of degrees of freedom, covariates/factors used, model details, programs implemented, etc. For example, knowing these aspects might easily clarify observed variability. Unfortunately, this information is still not provided in many cases, and hiding it makes meta-analyses difficult.

The above recommendations apply quite generally beyond voxelwise and region-wise brain studies. For example, connectograms can be shown with the highlighting methodology. Furthermore, these steps should be complemented by sharing the unthresholded results (both the statistics *and* the effect estimates, wherever possible) and analysis codes publicly, facilitating meta-analyses.

Many of these recommendations are in line with the philosophy of the current COBIDAS recommendations for data analysis and sharing (Nichols et al., 2017). They also expand on a couple of key points in ways that are important. Firstly, COBIDAS only refers to model validation in the context of machine learning analyses; instead, model validation can, and should, be applied whenever possible. Secondly, including effect estimates would often greatly expand the richness and information content of reported results (Chen et al., 2017); more results should include these values. Thirdly, we strongly agree with Nichols et al. (2017) about the central importance of figures in a paper:

> “*The thresholded map figures perhaps garner the most attention by readers and should be carefully described*.*”*^11^

We would just amend this statement to recommend the application of *transparent* thresholding in figures, for all the reasons detailed above.

### Cross-study comparisons and meta-analyses

During FMRI processing, there are many choices made and parameters set (e.g., see Carp, 2012). Many of these have what might be called “semi-arbitrary” ranges. For example, there is no clear, single answer as to whether one should smooth a 2mm isotropic EPI voxel by 0, 1, 2, 2.75, 3.14159, 4 or 5 mm when performing a voxelwise study (or perhaps by more, or whether to blur it within a brain or tissue mask, etc.), though there is likely consensus that smoothing by 20 mm would not be acceptable for human datasets.^12^ Each of these choices will affect the outcome in some way and perhaps reflects a slightly varied assumption, goal or simply “lab default”, but such a range or set of values seems reasonable---hence, the “semi-arbitrary” description. The reasonable range is based on an understanding of the processing step’s purpose; for smoothing, it might be the set of values that balance usefully blurring noise with preserving local structure, and the 20 mm blur would violate the latter property. In practice, choosing a reasonable parameter value from a semi-arbitrary interval is part of the “analytic flexibility” observed across studies; such choices occur at many stages of processing, and this does create an inherent aspect of variability, which likely cannot be avoided.

In the same way, there is also analytic flexibility in the choice of meta-analytic methods and cross-study comparisons. There is a large variety of possible approaches, which utilize different data content and very different assumptions. These are important to understand. Some meta-analysis methods are inherently sensitive to minor differences or tuning parameters, and therefore they tend towards reporting low similarity when the actual differences in results are small. For example, Method A might threshold data before comparison, and Method B might compare unthresholded results. Both methods depend on the data values, but Method A reduces the amount of data in the comparison and is also sensitive to the (arbitrary) threshold value. In FMRI in particular, thresholded boundaries themselves will depend strongly on features such as number of trials or subjects in a study, the exact statistical model used (orthogonalization procedure or parametric/amplitude modulation), etc. Method A is also in line with the interpretation of FMRI results representing ON/OFF brain responses, as well as essentially all of the other disadvantages of strict thresholding listed at the start of this section.

Therefore, it seems difficult to consider the differences in using meta-analysis Method A or B to be merely semi-arbitrary. They are likely to provide quite different results in many cases, which are sensitive to very different features within the data. The original NARPS paper reinforces this view, as it contained two similarity analyses across participating teams that reached very different conclusions from the exact same data:

1. When addressing Yes/No questions based on all-or-nothing thresholding (essentially Method A), there was relatively high variability among teams, summarized by the authors as, “sizeable variation in the results of hypothesis tests.” (Three hypotheses had 94% agreement across the teams; three hypotheses had about 80% agreement; and three hypotheses had approx. 65% agreement.)
2. When analyzing unthresholded results (essentially Method B), there was much lower variability among different teams; and furthermore, the variation was mainly in response magnitude, not sign. Indeed, the unthresholded teams’ submissions were “highly correlated.” This point is furthered by Lindquist (2020), endorsing the following suggestion based on NARPS findings:

> *“The first is to share unthresholded activity maps, because this will allow image-based meta-analysis. The authors found that such an analysis, which aggregated information across teams, yielded consensus results, no doubt aided by the fact that the spatial patterns in the activity maps were highly correlated across groups*.*”*

Indeed, these results suggest that the largest source of analytic variability in NARPS was not any processing step, but instead the choice of meta-analysis itself.

In general, using a meta-analysis method that most accurately represents the full content of the underlying data, and minimizes sensitivity to tuning parameters, seems desirable. This framework is consistent with the highlighting methodology, whose benefits have been noted above. If unthresholded results are important to share for meta-analyses, then they should be important enough to visualize in the main paper.

In the case of the NARPS analysis, such an approach (Method B or #2 above) also showed that the teams’ results generally had a high degree of reproducibility. Whether looking across the whole brain or zooming in on ROI-focused locations, the correlations of unthresholded statistics strongly suggested a great deal of similarity across the teams. The greatest amount of variability appeared to be due to a sign flip in some teams’ results, which was typically explainable and correctable (and clearly observed when viewing the full results, importantly). However, the results strongly suggest that pipeline variability across teams did not produce “sizeable variation” in a random or haphazard way. Instead, the majority of results showed very high agreement in both the *location* and *directionality* of effects, and differed mainly in statistical strength. It is true that if such results are compared after thresholding, there would be variation in the scientific conclusions. However, when viewed in the “highlighting” mode, most of those conclusions would strongly agree on the important aspects of the study---the major locations and directionality of the strongest effects---varying mainly in degree. The variability among NARPS teams primarily reflected deviation in the degree of agreement, rather than conflict among results. This is demonstrated in the original NARPS paper Fig. 2a, which is very similar to Fig 8B here (and their Ext. Data Fig. 2 corresponds with the matrices calculated here and presented in the Supplementary Info).

The neuroimaging community is rightfully concerned with the issue of reproducibility in the field, and about understanding the role that processing and analytic flexibility has. These should be understood. There is a degree of inherent variability due to the semi-arbitrary choices made, and perhaps some will be reduced over time after further investigations. This must be considered within the context of other sources of variability within the study, such as scanner (strength, vendor, software version, shimming, distortions, and more), acquisition parameters (voxel size, TR, etc.), study size, paradigm structure, and research hypotheses. Furthermore, as shown in the NARPS study and here, the style of meta-analysis plays a hugely important role in combining studies; comparing unbinarized results is preferable. Much of this has to do with included information content available for comparison. For example, peak voxel results contain much less information than volumetric comparisons, and (as demonstrated above) can be unstable and unrepresentative of results; in a review of neuroimaging meta-analyses, Wager et al. (2009) strongly preferred that:

> *“Alternatively, full statistic maps for each study could be used and effect sizes aggregated at each voxel (Lewis, 2006). Though we consider this to be a “gold standard” approach, and advocate its development in future meta-analytic work, this approach is complicated by the lack of readily-available statistic images*.*”*

Fortunately, modern data sharing abilities make this feasible (and should include effect estimate comparisons, as well).

### Conclusions

At its heart, data processing and analysis involves taking a large amount of data as input and bringing it to a state from which meaningful, generalizable conclusions can be made. The final stage of a study is to interpret the results, and to present them in an accurate way for others to understand. Providing more complete results reports helps both the study researchers and other readers make accurate interpretations---they can reduce the chance of a conclusion being “overfit” to a sparse set of outcomes. As we have seen in many examples here, information contained in sub-threshold locations is often extremely useful in understanding the supra-threshold ones, whether by providing context, an insight into deeper spatial patterns, quality control, or signs of particular differences. This must be balanced with not over-providing data, which can overwhelm or reduce interpretability, and the “highlighting” methodology aims to still avoid that.

No single neuroimaging study will definitively prove an effect or a hypothesis or an association, so we should avoid forcing studies to artificially act as if they must make a binarized decision about each element (ON/OFF, etc.). Understanding the activity and interaction of the hierarchically complex brain can only be done by an accumulation of work over many studies, each using different data collections, paradigms and methods to thoroughly explore a finding. Each study should be designed and reported in a way that facilitates comparisons and pooling of information---recall that statistics values themselves are descriptions, not decision-makers. Hiding away results, as in traditional FMRI results reporting, even if they have slightly lower statistical significance than the chosen threshold, inhibits this effort. The “highlighting” methodology promotes better science by sharing more information with the neuroimaging community from the very start. This is particularly true when the emphasis of study digestion lies with the figures and summary points drawn from them. We have seen several examples here of how different styles of reporting of the same results can lead to drastically different conclusions or ambiguity, and that all of them represent the full set of data at hand. As a scientific community, we should choose to utilize those that provide more accurate information.

The present work does not directly evaluate methods for cross-study comparison and reproducibility metrics explicitly, which are numerous and have various applications. That is an important but separate topic. However, the results here do suggest that primary comparisons and meta-analyses should favor those metrics that have minimal dependence on thresholding, which can distort perception of the data. For example, Dice coefficients are quite sensitive to thresholding, and the locations of peak statistics in clusters can be highly sensitive to minor processing choices and subject inclusion/exclusion criteria. Such methods treat individual studies as “decided results” to be compared rather than as pieces of evidence to be combined. Through the investigation here, and through the NARPS own results, the importance of the latter view has been demonstrated.

Data sharing initiatives are also important and should be used more readily across the field. Several aspects of the present investigation would not have been possible without them. However it must be emphasized that the initial impact and reception of a study lies with the results reporting. A much smaller fraction of the scientific community go through the steps to locate, download and re-investigate shared data than those who read a paper, view a conference poster or scan a summary. Even in science, “a picture is worth a thousand words”---or possibly more---so reporting results more fully is important.

## Acknowledgments

PAT, RCR and GC were supported by the NIMH Intramural Research Program (ZICMH002888) of the NIH/HHS, USA. VDC was supported by NSF grant #2112455. JG-C, DH and PAB were supported by the NIMH Intramural Research Program (ZIAMH002783) of the NIH/HHS, USA. AM was supported by a NIBIB grant (R01EB027119) of the NIH/HHS, USA. This work utilized the computational resources of the NIH HPC Biowulf cluster (http://hpc.nih.gov). We also thank the NARPS organizers and all of the team participants. While we emphasize different features than the main study, we also highlight points of agreement. We appreciate their work, their thoroughness and the clarity of their shared outputs, all of which greatly facilitated the examples presented here.

### Appendix A: Highlighting in AFNI

There are many ways to implement the highlighting visualization, which was first implemented in AFNI in 2014. The threshold-based transparency is determined by setting the “alpha” level within the RGBA color representation of the overlay, where alpha=1 is opaque, alpha=0 is perfectly transparent, and the intermediate values are translucent. The alpha-based opacity modulation can be activated within the AFNI GUI by clicking the “A” button in the overlay controller, or by setting an environment variable in the startup file. When activated, by default the opacity of a sub-threshold overlay voxel decreases quadratically from alpha=1 at the threshold value, to alpha=0 when the value is zero; the transparency can also be set to decrease linearly.

It is also possible to set the maximal opacity in the image to be <1, e.g., to be 70% opaque, so that underlay structures are visible everywhere. In such cases, sub-threshold translucency is applied in the same way as described above, just decreasing quadratically or linearly from the chosen maximum opacity value.

Separately, placing a line around the threshold boundary (placing a “box” around the region) is activated in the AFNI GUI either by clicking the overlay controller’s “B” button or by setting a separate environment variable. One could highlight results using either “A” or “B” separately, as well as together, depending on the application.

When the clusterizing functionality is activated in the GUI---that is, implementing a cluster-based threshold, in addition to a voxelwise one---the “A” and “B” functionality adapt to consider the additional threshold, as well. That is, only “large enough” clusters are opaque and outlined. Finally, we note that the AFNI program @chauffeur_afni, which provides a convenient command line and scripting interface for creating volumetric images and montages also contains options for including the transparency and outlining based on voxel thresholds and/or cluster thresholds. This command was used to generate most of the brain images in the present work.

## Supplementary Information

This supplementary section contains “highlighting”-style images of the results of the 61 NARPS teams for the remaining hypotheses (only data for Hyp #1 and 3 were shown in the main paper). For Figs. S1-6, panel A contains images of the transparently thresholded statistical datasets; see the Results and particularly Fig. 7-8 in the main paper for description details. Similarity matrices of both “hiding” and “highlighting” approaches are shown in panel B of each figure, using Dice coefficients and Pearson correlation, respectively. Following Botvinick-Nezer et al. (2020), the metadata responses by each team about whether the positive values in the image represent positive or negative activation have been applied, for consistent interpretations of Hyp #5, 6 and 9. In all cases, a great deal of similarity of the NARPS teams’ results is apparent for each hypothesis when viewing the informative “highlighting”-style images and matrices.

**Figure S1.**
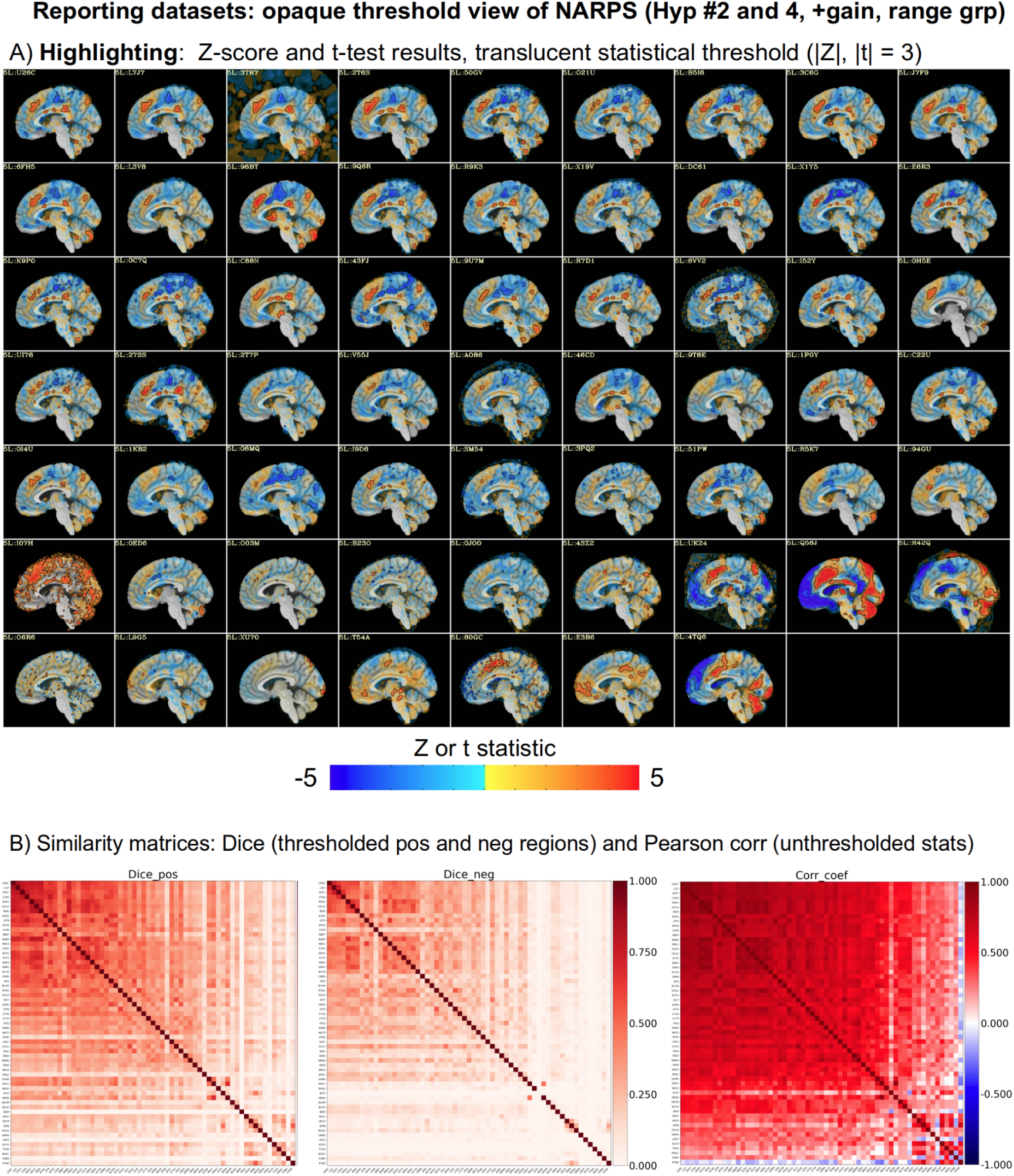
See Figs. 7-8 in the main paper for details. A) Transparently thresholded images across the original NARPS teams results for the positive parametric effect of gain in the equal range group (Hyp. #2 and 4). B) Similarity matrices describing the datasets in panel A: Dice coefficients (“hiding” summary) and Pearson correlation (“highlighting” summary). The vast majority patterns of positive and negative effects are quite similar across groups, while just the magnitudes of the statistics differ.

**Figure S2.**
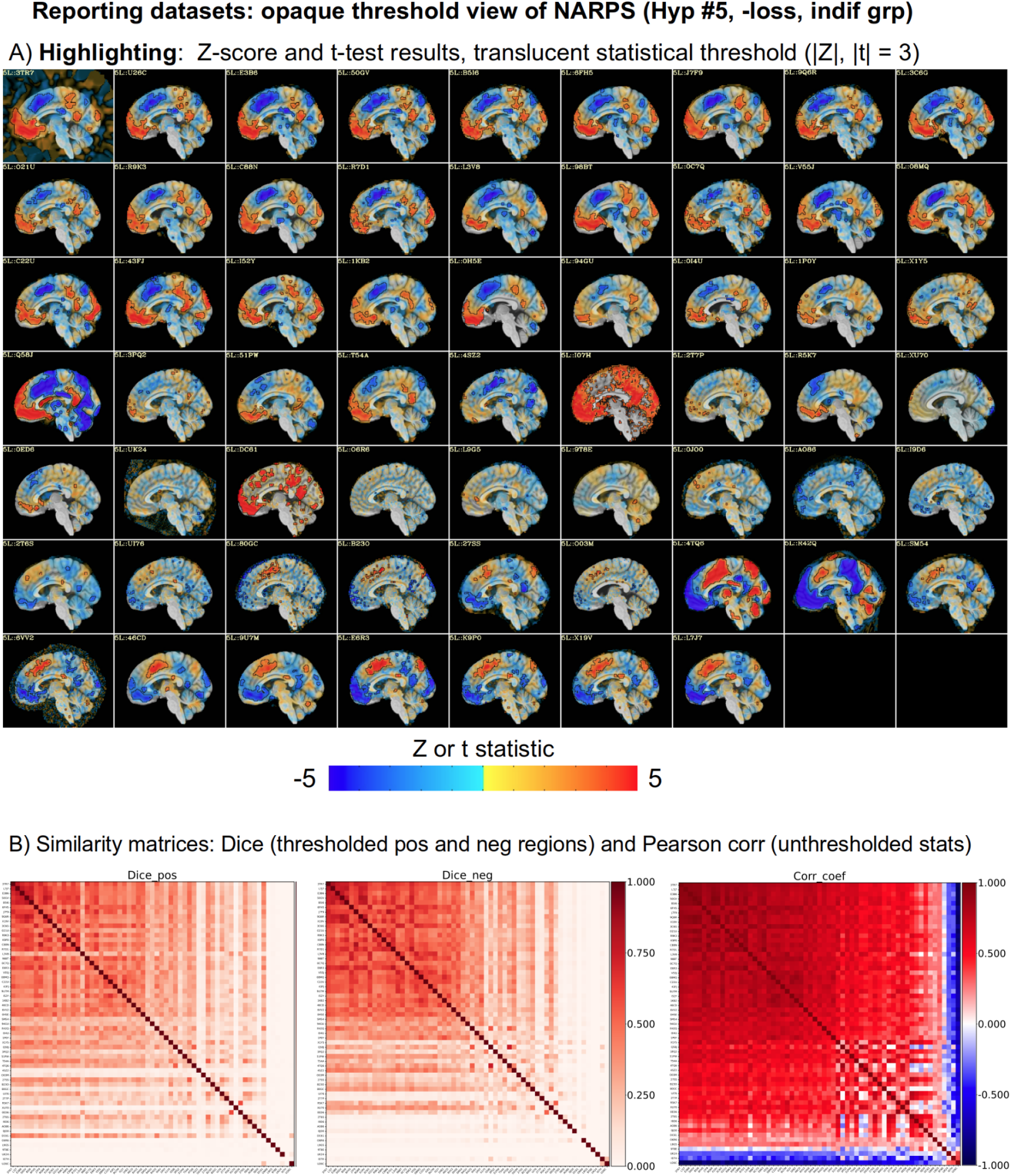
See Figs. 7-8 in the main paper for details. A) Transparently thresholded images across the original NARPS teams results for the negative parametric effect of loss in the equal indifference group (Hyp. #5). B) Similarity matrices describing the datasets in panel A: Dice coefficients (“hiding” summary) and Pearson correlation (“highlighting” summary). The vast majority patterns of positive and negative effects are quite similar across groups, while just the magnitudes of the statistics differ. A small number of teams appear to have a flipped sign of responses (strong anticorrelation; blue region in panel B’s Pearson correlation matrix).

**Figure S3.**
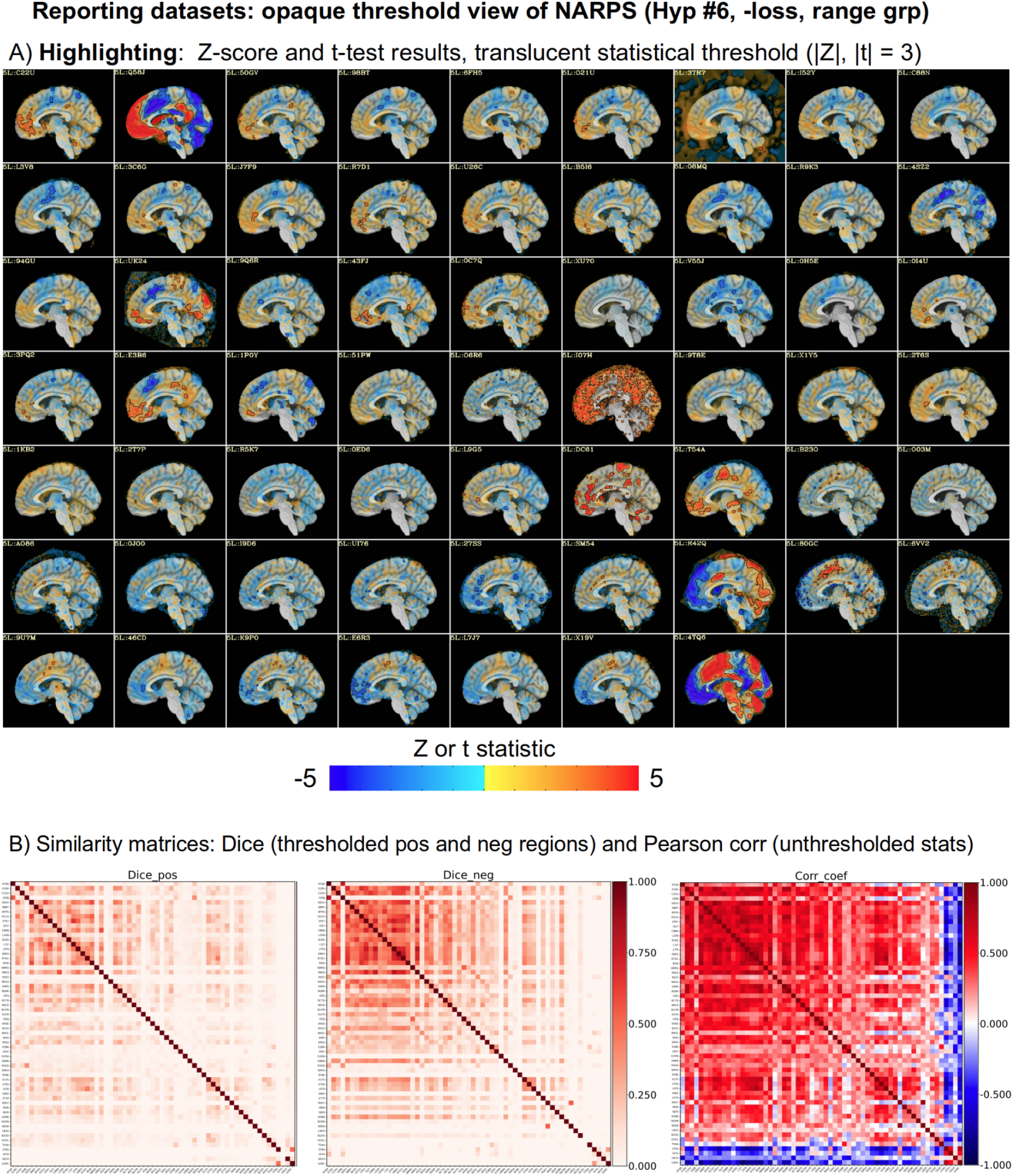
See Figs. 7-8 in the main paper for details. A) Transparently thresholded images across the original NARPS teams results for the negative parametric effect of loss in the equal range group (Hyp. #6). B) Similarity matrices describing the datasets in panel A: Dice coefficients (“hiding” summary) and Pearson correlation (“highlighting” summary). The vast majority patterns of positive and negative effects are quite similar across groups, while just the magnitudes of the statistics differ. A small number of teams appear to have a flipped sign of responses (strong anticorrelation; blue region in panel B’s Pearson correlation matrix).

**Figure S4.**
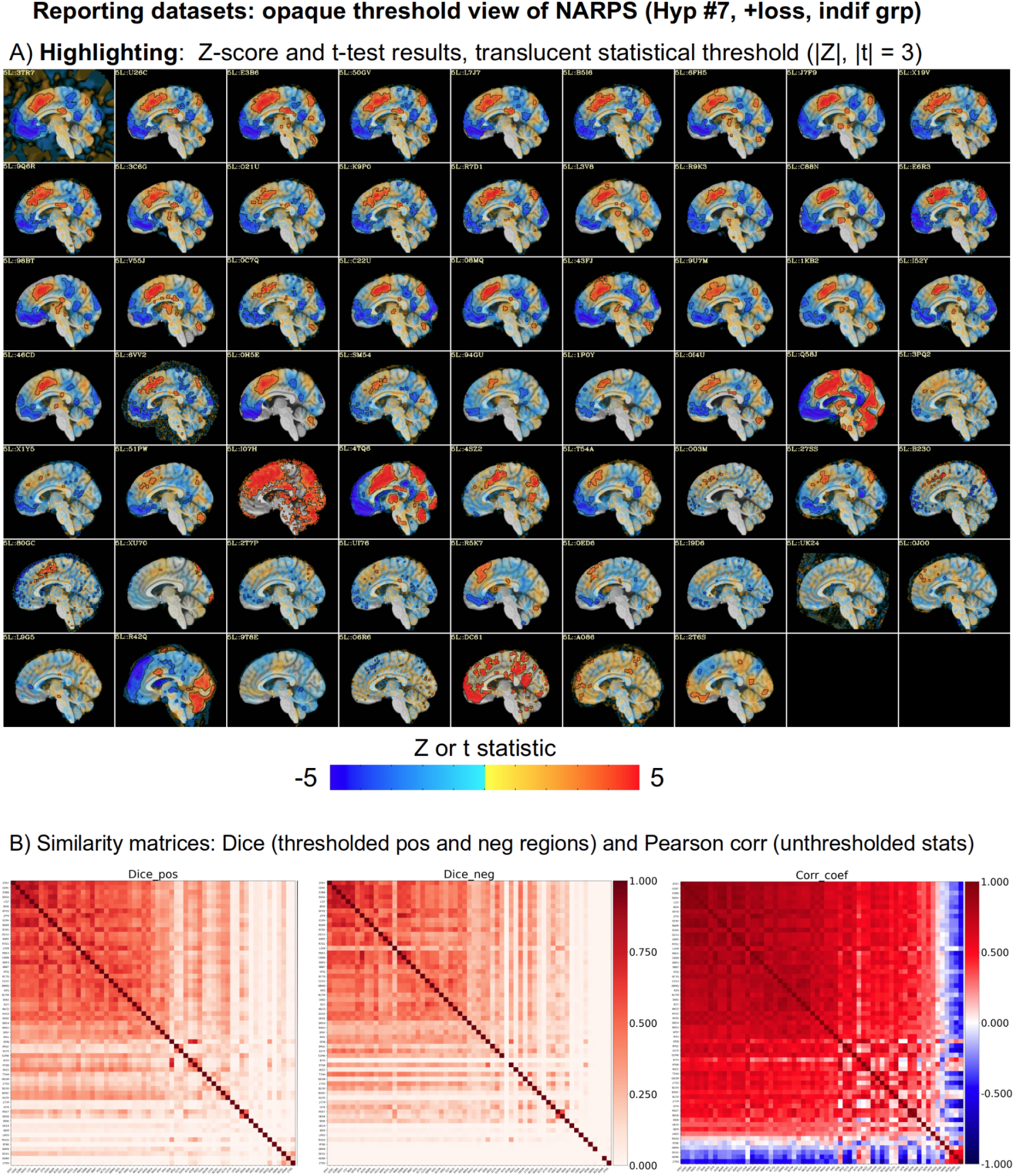
See Figs. 7-8 in the main paper for details. A) Transparently thresholded images across the original NARPS teams results for the positive parametric effect of loss in the equal indifference group (Hyp. #7). B) Similarity matrices describing the datasets in panel A: Dice coefficients (“hiding” summary) and Pearson correlation (“highlighting” summary). The vast majority patterns of positive and negative effects are quite similar across groups, while just the magnitudes of the statistics differ. A small number of teams appear to have a flipped sign of responses (strong anticorrelation; blue region in panel B’s Pearson correlation matrix).

**Figure S5.**
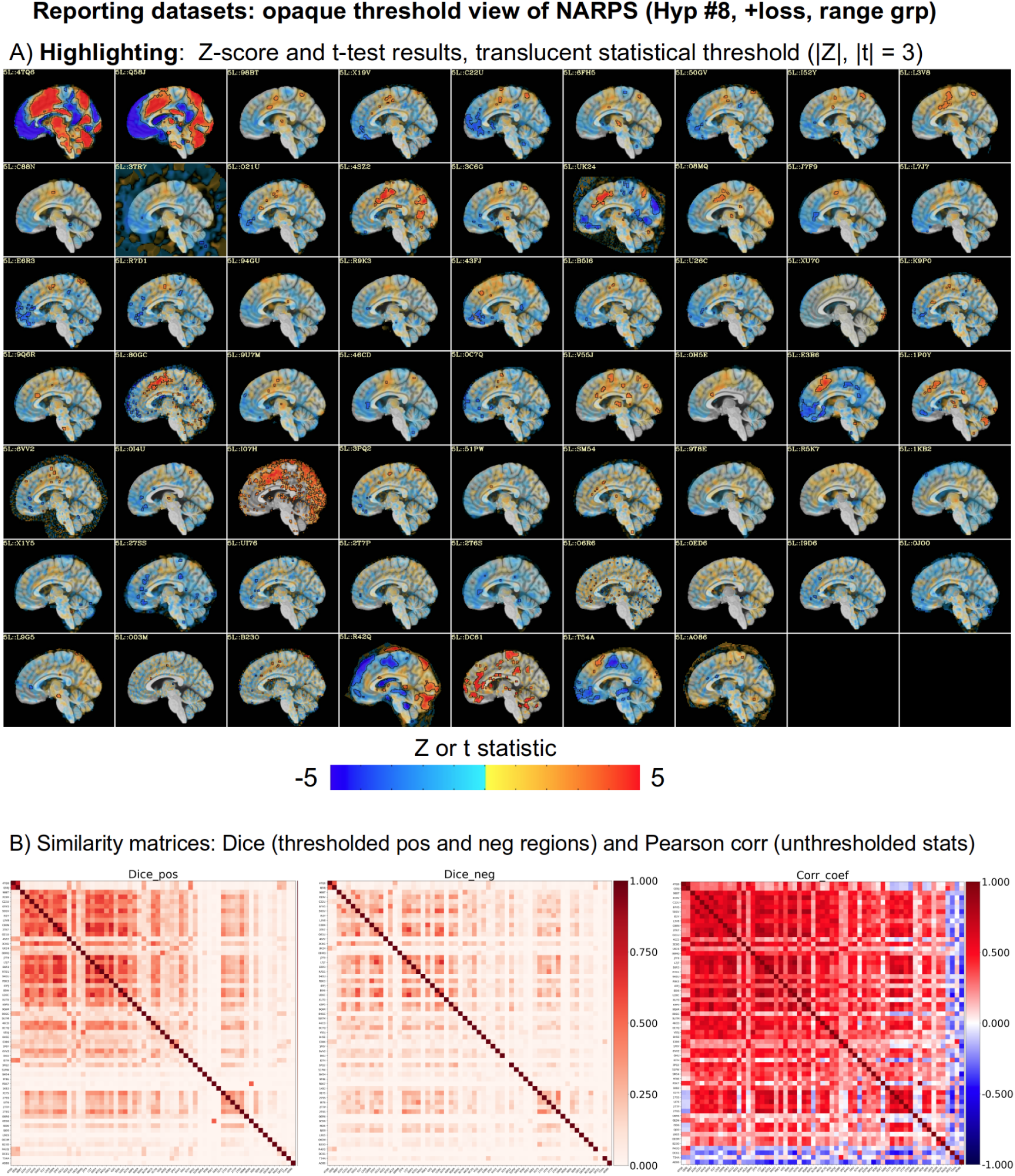
See Figs. 7-8 in the main paper for details. A) Transparently thresholded images across the original NARPS teams results for the positive parametric effect of loss in the equal range group (Hyp. #8). B) Similarity matrices describing the datasets in panel A: Dice coefficients (“hiding” summary) and Pearson correlation (“highlighting” summary). The vast majority patterns of positive and negative effects are quite similar across groups, while just the magnitudes of the statistics differ. A small number of teams appear to have a flipped sign of responses (strong anticorrelation; blue region in panel B’s Pearson correlation matrix).

**Figure S6.**
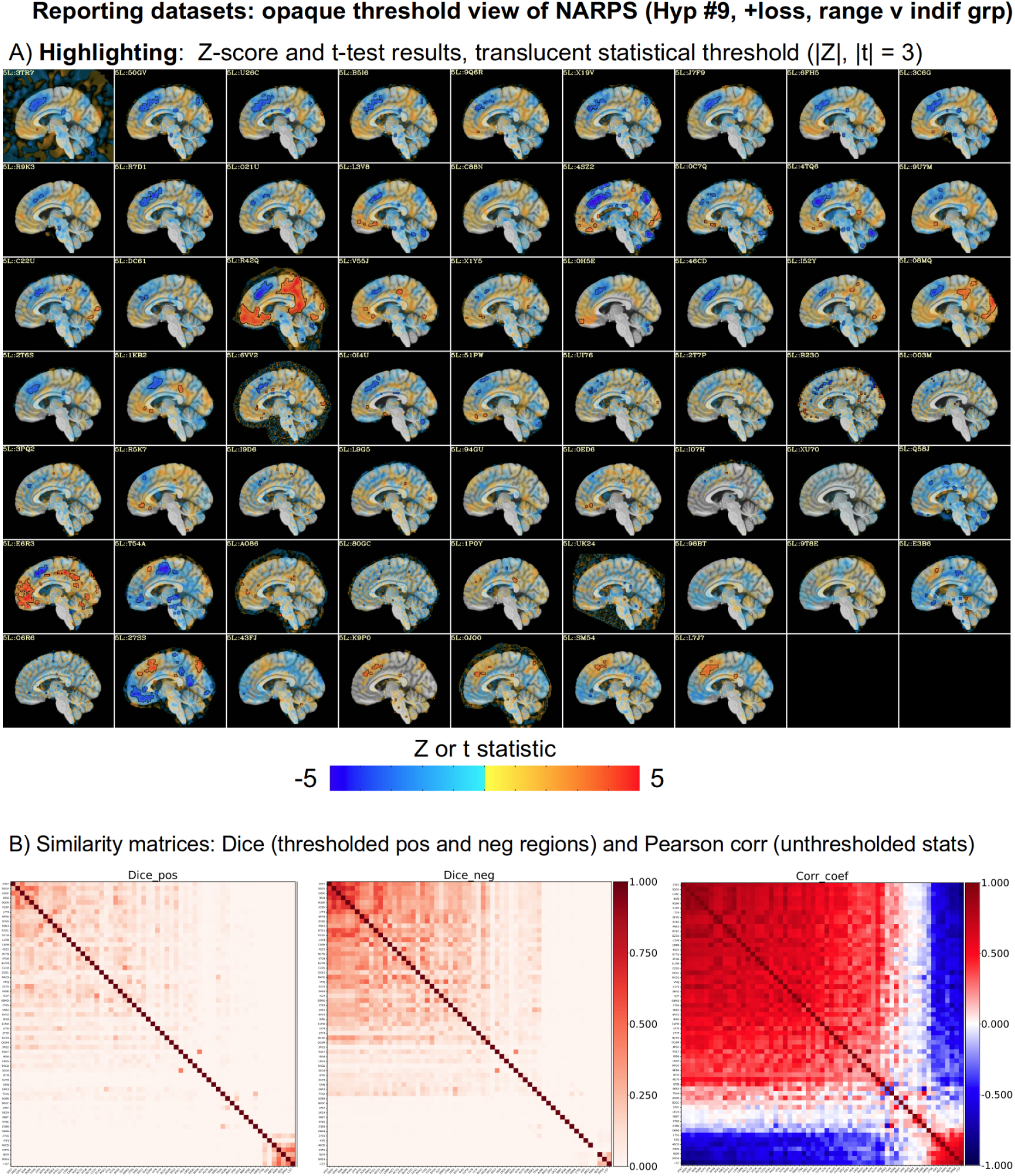
See Figs. 7-8 in the main paper for details. A) Transparently thresholded images across the original NARPS teams results for the positive parametric effect of loss in the equal range vs equal indifference groups (Hyp. #9). B) Similarity matrices describing the datasets in panel A: Dice coefficients (“hiding” summary) and Pearson correlation (“highlighting” summary). The vast majority patterns of positive and negative effects are quite similar across groups, while just the magnitudes of the statistics differ. A notable fraction of teams appear to have a flipped sign of responses (strong anticorrelation; blue region in panel B’s Pearson correlation matrix), and some have almost no correlation with the main block (white region in Pearson correlation matrix).

## Footnotes

1 Here, we focus on examples of whole brain FMRI and clustering. But the same problems with traditional thresholding apply in many other analyses, such as for region- and matrix-based similarity measures of correlation, coherence, mutual information, entropy, graph theory, etc. In many cases, subthreshold information is hidden away and the results depend sensitively on the essentially arbitrary threshold value(s). Therefore, in this discussion, references to “voxel” in example context generally apply more widely (to the scenarios with regions, matrix elements, etc.), since the same kinds of concerns arise.

2 There has been a transition towards uploading unthresholded data into online repositories linked to papers for other groups to investigate. This is a positive trend, but it does not fully ameliorate the negative consequences of hiding data within the main presentation in a paper, poster or other medium.

3 The Matlab implementation of this used by Allen et al. is available online: https://trendscenter.org/x/datavis/ This highlighting feature has been available in the AFNI software package since 2014, and Appendix A provides usage notes and details of how translucency is applied.

4 Hyp #1-8 were each 1-sided hypotheses for a single effect in a single group (e.g., Hyp #1: “positive effect [of gain] in VMPFC - for the equal indifference group”). Hyp #9 was a 1-sided hypothesis for a contrast between the two groups: “greater positive response to losses in amygdala” for the equal range group vs the equal indifference group.

5 After the end of the project, three teams changed responses about certain hypotheses. This agreement increased by one team for each of Hyp #3, 4 and 6, and decreased by two teams for #5.

6 Note that this model validation for the t-tests can only be performed through Bayesian modeling with the matched conventional assumptions, as the PPC is done through simulations. Since RBA is itself a Bayesian model, the PPC checks follow naturally.

7 Here, the same statistical information is used for both overlay coloration and thresholding, as effect estimate maps were generally not available).

8 As we have presented Fig. 7A, one might suspect a sign flip in some results, since we have presented both positive- and negative-effect clusters simultaneously. If only the former had been shown to match the stated 1-sided hypotheses, this would become even more difficult. This shows further benefit of the more informative highlighting methodology in understanding the presented data, even when it shows potential conflict of results.

9 The NARPS dataset, and its meta-analysis of group results that differ only in processing and analysis choices, is quite useful but is not a typical cross-study comparison framework. Even so, the highlighting methodology applies usefully to it, as presented above and discussed more below.

10 For similar reasons, we also suggest not masking data when processing, or even when presenting results, when possible. While not strictly a statistical process, it can also hide away signs of artifacts that would be better to know about. For examples of both using transparent thresholds and not masking during processing, see afni_proc.py-generated QC HTML reports.

11 COBIDAS Report: http://www.humanbrainmapping.org/COBIDASreport

12 Across participating NARPS teams, the reported range of applied smoothing for the 2 mm isotropic EPI voxels was 0-9 mm, with a median of 5 mm.

